# Neural Control of Autonomic Arousal During Threat Anticipation Revealed by High-Resolution Cardiac Contractility

**DOI:** 10.64898/2026.02.23.707545

**Authors:** Joanne E. Stasiak, Jingyi Wang, Neil M. Dundon, Elizabeth J. Rizor, Parker L. Barandon, Christina M. Villanueva, Viktoriya Babenko, Taylor Li, Kiana M. Sabugo, Scott T. Grafton, Regina C. Lapate

## Abstract

The sympathetic nervous system prepares the organism for adaptive action by shaping physiological, affective, and behavioral responses to environmental demands. Yet, how sympathetic signals dynamically couple with neural systems supporting emotional experience and behavior remains poorly understood, in part because common indices such as skin conductance responses lack sufficient temporal resolution to track these dynamics. Here, we evaluated trans-radial electrical bioimpedance velocimetry (TREV), a non-invasive measure of beat-to-beat cardiac contractility, and compared it with skin conductance responses during threat anticipation and simultaneous fMRI. Participants (n=60) completed a threat-of-shock paradigm requiring goal-directed action. Cardiac contractility increased during threat anticipation, covaried with skin conductance responses, and independently predicted self-reported emotional intensity. Critically, threat-related increases in contractility—but not skin conductance—tracked threat-related modulation of activation in dorsomedial prefrontal cortex, posterior parietal cortex, and cerebellum, with contractility-modulated cerebellar activation predicting faster motor responses under threat. These findings establish TREV-derived cardiac contractility as a physiological signal linking sympathetic drive with neural responding, emotional experience, and adaptive behavioral mobilization during emotion-guided action.

## Introduction

The autonomic nervous system (ANS) is central to the generation of adaptive responses to dynamic environmental demands—coordinating physiological adjustments across internal and external changes (Buijs, 2013; Porges, 1997). Emotions mobilize the body, including through sympathetic changes that prepare the organism for action (Adolphs & Andler, 2018; Scherer, 2014). Sympathetic nervous system (SNS) activation is especially critical for rapid mobilization in the face of threat, facilitating approach and avoidance-related behaviors through temporally precise modulation of cardiovascular and somatic systems (Jansen et al., 1995; Lang, 2014). Recently, a novel technique—trans-radial electrical bioimpedance velocimetry (TREV)—has been introduced as a noninvasive approach for capturing SNS activation with high temporal precision (Stump et al., 2024). Here, we situate TREV within established sympathetic indices, demonstrate its feasibility for simultaneous acquisition during fMRI, and underscore its sensitivity to subjective emotional experience and its associations with activation in brain regions linked to sympathetic drive under threat.

Decades of psychophysiological research employing common peripheral markers of SNS activity have established their utility and reliability for elucidating cognitive and emotional processes (Bradley & Lang, 2000; Braithwaite et al., 2015; Critchley, 2002; Crone et al., 2004; Kreibig, 2010). However, commonly used indices of sympathetic activity—such as electrodermal and impedance cardiography measurements—are constrained by several methodological limitations. For instance, skin conductance, a measure of eccrine sweat gland activity exclusively under sympathetic control (Dawson et al., 2007), is ubiquitous in psychological research and has been robustly linked to cognitive processing and emotional arousal (Bradley & Lang, 2000; Braithwaite et al., 2015; Kreibig, 2010). However, the relatively slow latency of skin conductance (e.g., 1–4 s to peak) requires long interstimulus intervals to reliably discriminate independent skin conductance responses (SCRs), reducing the temporal precision of the signal (Benedek & Kaernbach, 2010). Moreover, between 10–25% of individuals are typically classified as non-responders (Braithwaite et al., 2015; Dawson et al., 2007; Venables & Mitchell, 1996). Offering finer temporal specificity than skin conductance, impedance cardiography (ICG) is a well-established approach that uses the pre-ejection period (PEP) to index sympathetic influences on cardiac contractility (Bernstein, 2009; Woltjer et al., 1997). ICG can reliably be acquired in the MRI environment (Cieslak et al., 2015) and has been shown to be sensitive to emotional responding, cognitive load, and decision variables (Dundon et al., 2020, 2023; Kahle et al., 2016; Kreibig, 2010). However, ICG requires electrode placement over the thorax and neck, which renders this signal susceptible to respiratory noise; moreover, the dynamic range of PEP is narrow (Bernstein, 2009; Cieslak et al., 2018; Patterson, 1989; Sherwood et al., 1990; Stump et al., 2024). Together, these limitations highlight a critical gap in current psychophysiology toolkits: the lack of a temporally-precise sympathetic index that can be readily integrated with concurrent behavioral and neuroimaging paradigms.

A novel method for indexing cardiac contractility—TREV—addresses these limitations and holds particular promise for investigating psychophysiological processes in which dynamic autonomic changes are critical, such as in affective contexts that prepare an organism for action (Adolphs & Andler, 2018; Scherer, 2014; Stump et al., 2024). Offering distinct advantages over ICG and skin conductance, TREV provides a temporally precise and non-invasive index of sympathetic cardiac drive by deriving beat-to-beat estimates of ventricular contractility from impedance signals. Specifically, cardiac contractility is extracted from the second derivative (acceleration) of the arterial impedance waveform recorded at the forearm. Rapid changes in impedance reflect the pressure wave propagated through the arterial system at the moment of aortic valve opening during systole. The acceleration of this waveform indexes the magnitude of ventricular pressure generation during systole—a well-established proxy for myocardial contractility in a healthy heart and vascular system (Bernstein, 2009). As peak systolic pressure generation is directly modulated by sympathetic B-adrenergic input to the heart, these beat-to-beat impedance dynamics provide a physiologically grounded, non-invasive index of cardiac sympathetic drive (Stump et al., 2024). This computation obviates the need for concurrent ECG recordings and/or identification of subtle systolic landmarks in the impedance trace, such as the “B-point” required for PEP estimation (Kosicki et al., 1986; Stump et al., 2024). Collected via forearm electrodes placed over the radial and ulnar arteries (Stump et al., 2024), this configuration allows for quick participant preparation and eliminates susceptibility to thoracic and respiratory noise (Bernstein et al., 2012; Nagel et al., 1989; Stump et al., 2024). Prior work demonstrates that TREV-derived cardiac contractility is modulated by the anticipation of physical effort during a grip task (Stump et al., 2024), further validating TREV a reliable and high-resolution index of moment-to-moment fluctuations in sympathetic drive.

However, whether TREV is sensitive to experimentally-induced affective states, as well as whether it captures shared or distinct behavioral and neural correlates of sympathetic activity relative to commonly used SNS indices—such as skin conductance—remain unknown. Peripheral markers of SNS typically covary with the subjective arousal (or intensity) component of subjective emotional experience—but prior work has primarily been conducted using skin conductance responses or PEP-based indices of cardiac contractility (Cacioppo et al., 2000; Chatelain & Gendolla, 2015; Ganapathy et al., 2020; Herrald & Tomaka, 2002; Mendes et al., 2008; Sinha et al., 1992). As a result, it remains unclear whether TREV-derived cardiac contractility reliably captures self-reported emotional experience (such as subjective arousal) to an equivalent degree relative to other established SNS measures. Moreover, the feasibility of acquiring TREV concurrently with fMRI has yet to be demonstrated. Together, these gaps raise the question: does TREV index emotional arousal through the same neural circuits implicated in other sympathetic indices, or does it reflect partially distinct pathways of autonomic control?

On the one hand, sympathetic effectors share common subcortical neural architectures, with cardiovascular and sudomotor outputs arising from coordinated hypothalamic-brainstem pathways that converge on sympathetic preganglionic neurons (Critchley et al., 2011; Dampney, 1994; Dampney, 2016). However, these sympathetic response systems differ in their peripheral implementation and cortical regulation: whereas SCRs originate from acetylcholine-sensitive sudomotor neurons, cardiac contractility is predominantly governed by norepinephrine release from sympathetic postganglionic fibers (Cersosimo & Benarroch, 2013; Karemaker, 2017; Morrison, 1999; Wehrwein et al., 2016; Ziemssen & Siepmann, 2019). Consistent with this distinction, human neuroimaging studies indicate partially dissociable cortical control: lateral prefrontal regions have been more often associated with cardiac drive, whereas rostral and dorsomedial prefrontal regions have often been shown to reliably track electrodermal output, with shared engagement of subcortical regions such as the anterior cingulate cortex, hypothalamus, and amygdala during threat (Critchley, 2002; Dalton et al., 2005; Eisenbarth et al., 2016; Gianaros & Wager, 2015; Henderson, James, et al., 2012; Henderson & Macefield, 2013; Henderson, Stathis, et al., 2012). Of note, tract-tracing work in nonhuman primates demonstrates that motor and prefrontal regions with established roles in action selection and affect regulation directly influence sympathetic outflow, highlighting common integration of volitional and emotional control in autonomic responses (Dum et al., 2016). This top-down cortical influence provides a plausible mechanism through which different sympathetic effectors may be selectively engaged depending on task demands and emotional context. Together, these bodies of work suggest that different peripheral sympathetic measures may index overlapping yet partially distinct neural systems, further motivating examination of whether TREV is sensitive to affective contexts and whether TREV-derived cardiac contractility maps onto established circuits of autonomic arousal or reveals dissociable neural signatures.

Here, we simultaneously acquired TREV-derived cardiac contractility, skin conductance responses, self-reported subjective arousal ratings, and fMRI during a naturalistic threat-of-shock paradigm. Across an extended countdown anticipating shock, we orthogonally manipulated threat unpleasantness (mild vs. unpleasant shocks) and threat controllability (controllable vs. uncontrollable shocks). First, we examined whether both TREV-derived contractility and SCRs were reliably modulated by prolonged threat anticipation. Next, we evaluated convergent validity by testing whether the two sympathetic markers covaried with one another other. Building on prior evidence that skin conductance responses relate to the intensity of self-reported emotional experience (e.g. (Kreibig, 2010), we tested whether TREV-derived contractility was associated with subjective emotional intensity—and whether TREV explained unique variance in subjective arousal beyond SCRs. Finally, concurrent fMRI acquisition allowed us to examine neural correlates of TREV-derived contractility and SCRs and determine whether TREV captured distinct or overlapping neural signatures of sympathetic cardiac control. Together, these analyses provide the first simultaneous assessment of TREV, SCRs, and fMRI, offering critical new validation of TREV as a high-resolution tool for assessing sympathetic drive in cognitive and affective neuroscience research.

## Methods

### Participants

Sixty-nine participants were recruited at the University of California, Santa Barbara (UC Santa Barbara) (*M*_age_ = 20.59 years; *SD*_age_ = 2.00; range = 18–34 years; 51 F). Nine participants were excluded due to excessive motion (> 1mm mean framewise displacement across the functional run) and/or physiological artifacts (Skin conductance: undetectable SCRs in > 10% of trials; TREV: unidentifiable peaks in > 35% of each run’s timeseries), resulting on the final sample to comprise 60 participants (*M*_age_ = 20.52 years; *SD*_age_ = 1.89; range = 18–30 years; 47 F). Nine participants were classified as electrodermal non-responders (Benedek & Kaernbach, 2010); of those nine, six had low-quality TREV recordings and four exceeded motion thresholds. All participants were healthy, with normal or corrected-to-normal vision, and no self-reported history of psychiatric or neurological disorders. Written informed consent was obtained from every participant. All study procedures were approved by the UC Santa Barbara Human Subjects Committee. Participants were compensated monetarily for their participation.

### Procedure

Following informed consent and MRI safety screening, physiological sensors were applied to participants and electrical shock calibration was conducted. The Active Escape task was completed during ∼60 minutes of fMRI data collection. Upon completion, participants were paid for their time.

### Experimental design and statistical analysis

#### Active Escape Task

While undergoing fMRI, participants performed an Active Escape paradigm (adapted from Hur et al., 2020; see Stasiak et al., 2025 for details), consisting of seven functional runs (∼8 minutes each). Each run included twelve trials, equally divided between “Controllable Unpleasant”, “Uncontrollable Unpleasant”, “Controllable Mild”, and “Uncontrollable Mild” trials. Trial types were pseudorandomized such that no condition repeated consecutively (**Fig. 1a**). Each trial began with a 500ms fixation cross, followed by a 1000ms cue screen. The cue consisted of a background color signaling threat unpleasantness (red = Unpleasant shock, blue = Mild shock) and a letter indicating threat controllability (“O” = Controllable shock, “X” = Uncontrollable shock). After the cue, an 18-second countdown was presented, with each second marked by a decrementing number (18, 17, 16…). To minimize memory demands, the trial type (e.g., ‘Controllable Unpleasant’) was displayed as small text on the right side of the screen during the countdown. At the end of the countdown, participants were prompted to make a motor response using a handheld joystick, guiding a central cursor toward a target circle that appeared randomly in one of the four screen corners. Participants had 940ms to reach the target. In Controllable trials, successful motor responses (i.e., reaching the target circle within the allotted time) responses prevented shock delivery; in Uncontrollable trials, shocks were delivered regardless of response accuracy. Participants were instructed to respond on every trial.

**Figure 1.**
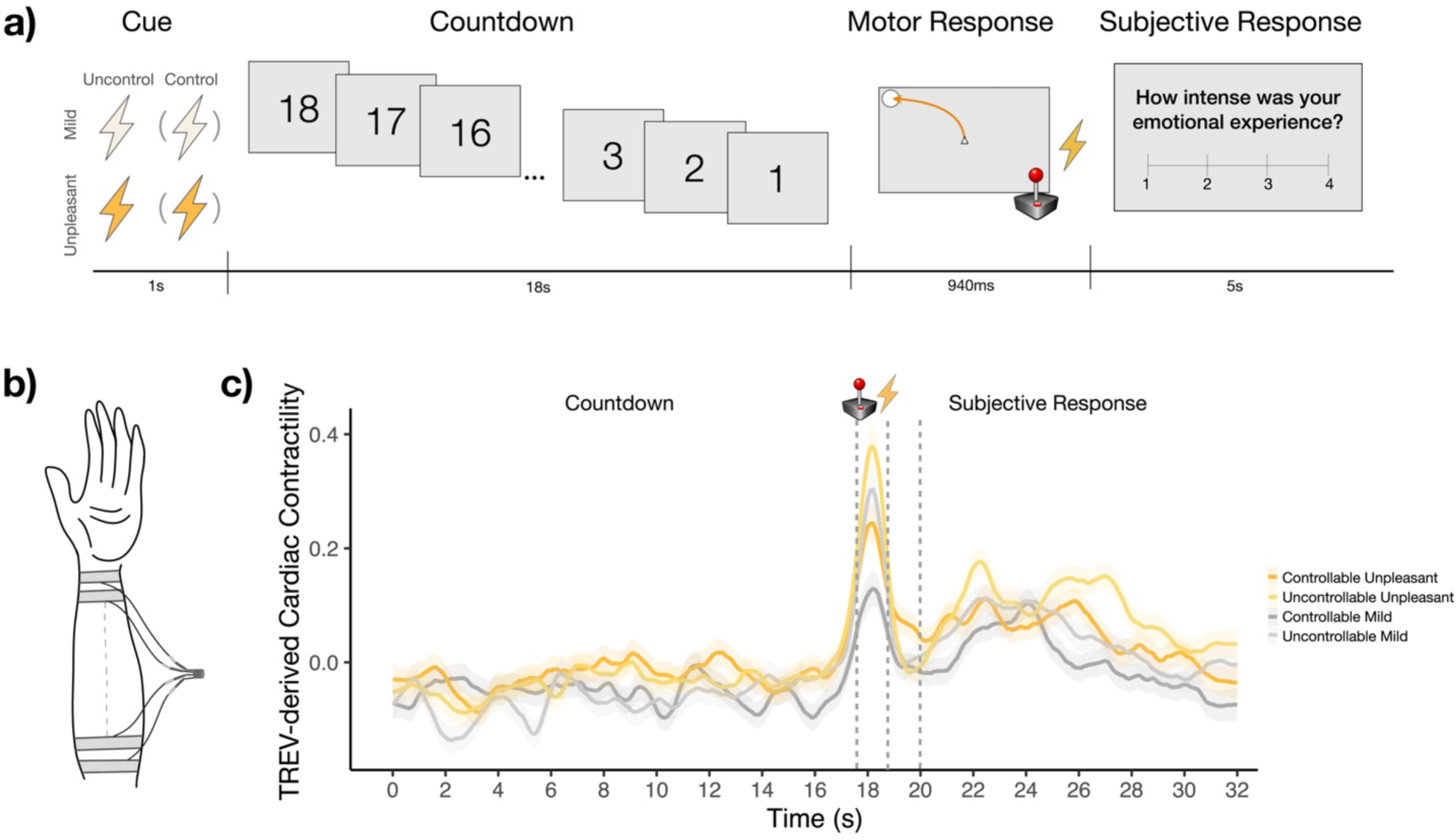
The Active Escape Task. **a)** In this threat-of-shock paradigm, participants tracked an 18s countdown to Mild or Unpleasant shock, which was manipulated orthogonally in relation to threat controllability (Controllable *vs*. Uncontrollable). In Controllable trials, shocks could be avoided by making a successful motor response, whereas shocks were administered regardless of performance in Uncontrollable trials. The motor response required using a joystick to make a time-sensitive (940ms), precise movement of a cursor to a target circle which was placed randomly in one of the four corners of the screen. At the start of each trial, participants saw a cue that informed them of the trial type (Controllable-Unpleasant, Controllable-Mild, Uncontrollable-Unpleasant, or Uncontrollable-Mild). Following each trial, participants reported the intensity of their emotions experienced during the countdown. **b)** Diagram of the TREV four-strip electrode placement on the forearm (adapted from Stump et al., 2024). **c)** The average timeseries of high-resolution cardiac contractility, measured via TREV, is plotted for the four experimental conditions.

Following conditional shock administration, a fixation cross was presented for 2000ms, followed by two questions assessing participants’ emotional experience. First, they evaluated the intensity of their emotional experience during the countdown (1 = *Not at all* to 4 = *Extremely intense*). Next, they rated their certainty in their emotional report (1 = *Not at all certain* to 4 = *Completely certain*; certainty ratings not analyzed here). Each rating was self-paced with a maximum of 5000ms. A variable intertrial interval (4000–6000ms) concluded the trial. The Active Escape task totaled 84 trials divided into 7 blocks and took ∼60 minutes to complete.

#### Shock Stimuli

Electrical stimulation was individually calibrated to each participant’s tolerance level, following procedures adapted from Hur et al. (2020). Initial calibration occurred while participants were positioned on the scanner bed but outside the magnet bore. Shocks were generated using an MRI-compatible constant-voltage stimulator (STM100C; Biopac Systems) and delivered through MRI-compatible electrodes (EL509; Biopac Systems) affixed to the inner surface of the right wrist. *<u>Mild calibration</u>*: Participants received a low-intensity shock (2.5 V) and were asked whether the shock was “reliably detectable” and whether it was “at all unpleasant” (Hur et al., 2020). If the shock was not reliably detectable, intensity was increased in 0.5 V increments until detectability was confirmed. If the stimulus was unpleasant, the intensity was decreased in 0.5 V steps until it was judged as clearly detectable but not unpleasant. This titrated value defined the “Mild” shock level used during the task (*M* = 21.24 V, *SD* = 4.71 V, range = 10–35 V). *<u>Unpleasant shock calibration:</u>* Participants received a 26.25 V shock and were asked whether the shock was as “unpleasant as they were willing to tolerate” (Hur et al., 2020). If not, shock intensity increased by 2.5 V and the question was repeated. This process continued until participants reported that the shock had reached their maximum tolerable level of unpleasantness. This value served as the “Unpleasant” stimulation during the task (*M* = 44.60 V, *SD* = 17.42 V, range = 26.25–103.75 V). A brief recalibration was completed inside the scanner bore to confirm that the previously established Mild and Unpleasant intensities remained appropriate.

#### Active Escape behavioral metrics

To examine whether sympathetic drive was associated with participants’ performance and experience in the task, we collected trial-wise measures of movement time and subjective emotional intensity. Movement time was operationalized as the time in seconds taken to successfully reach the target circle during the motor response period. Emotional intensity was measured by the subjective reports (*How intense was your emotional experience?* 1 = *Not at all* to 4 = *Extremely intense*) provided at the end of every trial.

### Physiological Data Acquisition

To measure sympathetic responding, we collected continuous recordings of EDA and TREV-derived cardiac contractility via Biopac System’s Acqknowledge 4.0 package. Prior to electrode placement, participants were instructed to wash their hands and left forearm with soap and water, then thoroughly rinsing with only water before fully drying.

#### Skin Conductance

Isotonic skin conductance gel was applied to one skin conductance electrode (Biopac EL501), which was placed on the participants’ left index finger and secured with medical tape. Data was collected with an MRI-compatible amplifier sampling at 2000Hz (EDA100C-MRI; Biopac Systems).

#### TREV

Researchers measured the circumference of participants’ left wrist before applying TREV electrodes, which were trimmed slightly if the 16cm electrode length exceeded participants’ wrist size to prevent electromagnetic loops in the MR environment. Four bioimpedance strip electrodes (BIOPAC EL526) were placed on the forearm (two electrodes on the distal region of the forearm and two electrodes on the proximal region of the forearm; see Stump et al., 2024, **Fig. 1b**). Proximal electrodes were placed approximately 1cm away from the wrist, and the distal electrodes were placed approximately 1 cm away from the elbow crease, with paired electrodes spaced about 1 cm apart. One TREV electrode served as a ground electrode that accounted for skin conductance’s required grounding. Data was collected with an MRI-compatible amplifier sampling at 2000Hz (NICO100C-MRI; Biopac Systems). Physiological data quality was assessed prior to the start of neuroimaging by instructing participants to take a series of deep breaths; subsequent waveforms were visually inspected prior to full data collection.

### Physiological Data Preprocessing

#### Skin Conductance

Skin conductance data was down-sampled to 10Hz and was analyzed in Matlab using Ledalab (http://www.ledalab.de). We used Continuous Decomposition Analysis (Benedek & Kaernbach, 2010), which deconvolves the SCR signal into phasic and tonic drivers of the skin conductance response. A detectable, stimulus-evoked SCR was defined as having a 1–4 second response window following stimuli onset and a minimum amplitude threshold of 0.02 µS (Boucsein et al., 2012). Participants without detectable SCRs in a minimum of 10% of the trials were classified as non-responders and were excluded from further analysis (n = 9 among the original n = 69 participants) (Benedek & Kaernbach, 2010). To correct for the skewed distribution of the skin conductance data, we square-root transformed SCR values(Boucsein et al., 2012).

#### Cardiac Contractility

TREV signals were processed in a Python-based signal processing software (Semi-automated Contractility estimates from Ohmic impedance measures within TREV (SCOT): https://github.com/caitg%20%20regor%20y/SCOT; cf. Stump et al., 2024). Within SCOT, a low-pass filter of 10Hz was applied to remove cyclical noise from the MRI gradient coils. All data were visually inspected to assess signal quality. For each participant, peak detection was applied to impedance acceleration waveforms to identify heartbeat event timepoints. Heartbeat timepoint markers were manually adjusted when software peak detection was found to be incorrect. Dubious peaks (such as those affected by motion or blunted by MRI signal noise) were replaced with the magnitude of the two preceding and two proceeding peaks by a trained researcher blinded to trial condition order and to how task cues (i.e., cue, countdown, response) aligned with cardiac data. Contractility waveforms were then computed by taking the derivative of the acceleration timeseries, which reflect the rate of ventricular pressure generation (Bernstein et al., 2012; Stump et al., 2024). Peak magnitude was computed for each contractility peak (whereby greater magnitude reflects greater contractility for that heartbeat) following automatic peak detection in SCOT. Respiration (derived from the low-frequency variance of the impedance velocity waveforms) and heart rate effects were modeled out of beat-wise contractility estimate data. Data were excluded from analyses if R-peaks could not be identified in ∼35% or more of the acceleration timeseries (n = 6 among the original n = 69 participants). Contractility values were z-scored for each participant prior to running additional analyses. For average timeseries of TREV-derived cardiac contractility across the four experimental conditions, see **Fig. 1c**.

### Physiology-Behavior Analyses

#### Active Escape task conditions

We examined whether TREV-derived contractility and SCRs were modulated by threat unpleasantness and controllability using linear mixed-effects models (with participant entered as a random effect) in R (lme4 package: version 1.1-38, RRID: SCR_015654; Bates et al., 2015). Anova (stats package: version 3.6.2, RRID: SCR_025678; Core Team, 2016) and emmeans (version 3.6.2, RRID: SCR_018734; https://github.com/rvlenth/emmeans) packages were used for follow-up contrasts. Summary metrics of sympathetic activity were computed by averaging TREV-derived contractility and SCRs across each trial’s countdown period; these aggregated trial-wise values for TREV and SCRs were used in subsequent models below.

#### Subjective Emotion

To investigate whether TREV-derived cardiac contractility and SCRs were associated with reported emotional intensity, we ran linear mixed-effects models (with participant entered as a random effect and threat unpleasantness and controllability entered as interacting variables) in R (lme4 package; Bates et al., 2015). In all models, emotional intensity was entered as the independent variable, while TREV-derived cardiac contractility and SCRs were entered as dependent variables in their respective models. To account for potential shared variance between TREV-derived contractility and SCRs, an additional model controlled for both sympathetic metrics (i.e., TREV and SCRs entered as simultaneous regressors).

#### Movement Time

To examine whether TREV-derived cardiac contractility and SCRs were associated with motor responding, we ran linear mixed-effects models (with participant entered as a random effect and threat unpleasantness and controllability entered as interacting variables) in R (lme4 package; Bates et al., 2015). For follow-up across-subject models, we aggregated across runs to obtain an estimate per condition and subject. These models were run using repeated-measures anova and linear regression models in R (stats package; Core Team, 2016).

### Functional MRI Methods

#### Image acquisition

Neuroimaging data were acquired in the UC Santa Barbara Brain Imaging Center with a Siemens Prisma 3T MRI scanner equipped with a 64-channel radio frequency head coil. Whole-brain Blood Oxygenation Level-Dependent (BOLD) fMRI data were obtained using a T2*-weighted 2 accelerated multiband echoplanar imaging sequence (54 axial slices, 2.5mm3 isotropic voxels; 80x80 matrix; TR = 1900ms; TE = 30ms; flip angle = 65°; 250 image volumes/run). High-resolution anatomical scans were acquired using a T1-weighted magnetization-prepared rapid gradient echo (MPRAGE) sequence at the beginning of the session for spatial normalization (TR = 2500ms; TE = 2.22ms; flip angle = 7°), followed by a gradient echo field map (TR = 758ms; TE1 = 4.92ms; TE2 = 7.38ms; flip angle = 60°).

#### fMRI data preprocessing

Functional neuroimaging data were preprocessed using FEAT (FMRI Expert Analysis Tool) Version 6.00, implemented in FMRIB’s Software Library(Jenkinson et al., 2012; Smith et al., 2004). Preprocessing steps included applying high-pass temporal filtering with a 100s cutoff, inclusion of standard and extended motion correction regressors obtained using MCFLIRT, spatial smoothing using a Gaussian kernel of 5mm, slice-time correction, and adding regressors of non-interest created from a matrix of points representing displacement of 0.5 mm or greater to control for movement-confounded activation. Brain extraction was carried out through Advanced Normalization Tools (ANTs) skull-stripping algorithm (Avants et al., 2009). Functional images were first registered to participant’s T1-weighted anatomical image using a linear rigid body (6-DOF) transform while maintaining native functional resolution (2.5 mm^3^ isotropic) and were then registered to standard space (i.e., the Montreal Neurological Institute (MNI) standard brain) using a linear affine (12-DOF) transformation in FNIRT. Following registration of functional masks to MNI-space, activation located at coordinates z < -40 was discarded due to cut-off cerebellar anatomy in the functional images.

#### fMRI data processing

Overview. We conducted whole-brain voxelwise regressions to examine whether threat-related changes in sympathetic drive, subjective emotional experience, and/or motor responding were associated with BOLD responding during anticipatory threat. Follow-up ROI analyses examined whether individual differences in contractility-related activation were associated with motor performance under threat.

#### Whole-brain univariate analyses

We used FSL’s three-level modeling approach (Jenkinson et al., 2012). In FEAT (FMRI Expert Analysis Tool, version 6.0.6, RRID: SCR_024915; Jenkinson et al., 2012; Smith et al., 2004), we first obtained run-wise BOLD activation parameter estimates using general linear models with a double-gamma hemodynamic response function. These models including eight regressors of interest, consisting of separate ‘countdown’ and ‘motor’ regressors for each trial type (Controllable Unpleasant/Controllable Mild/Uncontrollable Unpleasant/Uncontrollable Mild), in addition to one regressor of noninterest accounting for the subjective question-response periods (Stasiak et al., 2025). Then, a fixed-effects general linear model was computed to combine the parameter estimates across the 7 functional runs for each participant. Next, whole-brain associations were examined at the group-level using a random-effects models (FLAME). Automatic outlier de-weighting was run on a voxelwise basis. Activations were cluster-corrected for multiple comparisons across the whole brain at *Z* > 2.3, *p* < 0.05. Univariate results for this task (independent of sympathetic modulation or subjective experience) have been previously reported (see Stasiak et al., 2025).

### Parametric modulation

#### Sympathetic Metrics

To examine the distinct and common neural bases of changes in sympathetic drive as indexed by TREV and skin conductance, we ran two parametric modulation models of trial-wise BOLD responses using z-scored indices of TREV-derived cardiac contractility and SCRs, respectively (Heinzel et al., 2005). Separate parametric modulation models were constructed in which each sympathetic metric served as the trial-wise regression ‘weight’ (Büchel et al., 1998; Heinzel et al., 2005). In addition to the regressors described above, two regressor variables with sympathetic modulation parameters for the Unpleasant and Mild shock trials were included in these models. Because we did not observe significant effects of Controllability on either TREV-derived cardiac contractility or SCRs, our final models only included regressors for Unpleasant and Mild trials. To examine condition-independent sympathetic-related neural activation (i.e., positive and negative correlations with sympathetic activity regardless of task condition), an additional parameter estimate was created from the mean of the TREV-modulated Unpleasant and Mild regressors. The contrasts of interest in each model therefore included 1) Δ Unpleasant − Mild, 2) Positive Correlation, 3) Negative Correlation. We ran these models using the three-level process in FEAT (Jenkinson et al., 2012; Smith et al., 2004), as described above. Following whole-brain group-level analyses, we examined whether overlapping activations were observed across TREV-derived cardiac contractility and SCRs parametric modulation maps using a conjunction approach. To this end, activation maps of each contrast were first binarized, then multiplied across sympathetic channels (e.g., PositiveCorrelation_SCR * PositiveCorrelation_TREV), such that resultant maps would indicate clusters of activation present in both model contrasts.

#### Subjective Emotion

To examine potential overlap between sympathetic-derived BOLD modulated changes and BOLD changes associated with self-reported emotional arousal or intensity, we ran a model of neural activation parametrically modulated by trial-wise emotional intensity ratings. To this end, each trial’s emotional intensity rating was demeaned and included it as the trial-wise regression weight. Two regressor variables with emotion-modulation parameters for the Unpleasant and Mild shock trials were included in this model, where the primary contrast of interest was Δ Unpleasant − Mild. We ran this model using the three-level process in FEAT (Jenkinson et al., 2012; Smith et al., 2004), as described above. Following whole-brain group-level analyses, we tested potential common activations across subjective emotional intensity with both TREV-derived cardiac contractility and SCRs by multiplying the binarized activation maps (Δ Unpleasant − Mild) with the corresponding map for each sympathetic channel (e.g., Δ Unpleasant − Mild_Emotion * Δ Unpleasant − Mild TREV), whereby resulting maps indicate clusters of activation reliably present in both contrasts. Cluster-correction for multiple corrections was performed at *Z* > 2.3, *p* < 0.05.

To test for potential specificity in results observed in the results from parametric modulation models obtained for TREV relative to skin conductance (Δ Unpleasant − Mild), we conducted one-sample t-tests. We first subtracted beta weights from the corresponding activation maps for each participant (e.g., [Δ Unpleasant − Mild TREV] − [Δ Unpleasant − Mild SCRs]). Then, we tested this difference map against 0 using nonparametric permutation testing in FSL, which was corrected by multiple comparisons using threshold-free cluster enhancement (TFCE) (randomise; n = 5000 permutations (Winkler et al., 2014).

#### Region-of-interest analyses

Following our observation of significant contractility-related cerebellar activation under threat and given a breadth of research relating cerebellar activation to motor control (Dum & Strick, 2003; Ebner & Pasalar, 2008; Houk et al., 1997), we ran ROI analyses to investigate potential behavioral correlates of contractility-cerebellar interactions. Cerebellar ROIs were defined using the SUIT atlas (Diedrichsen & Zotow, 2015), thresholded at 50%, and registered from MNI to participants’ structural space using FNIRT (10 mm warp) while maintaining native resolution (2.5 mm^3^ isotropic).

## Results

### Sympathetic channels, emotional experience, and task performance are robustly modulated by anticipatory threat

We first verified that our threat-of-shock paradigm effectively induced changes in sympathetic drive, subjective emotional intensity, and goal-dependent task performance. Threat unpleasantness robustly increased TREV-derived cardiac contractility (*F* = 8.166, *p* = 0.004) (**Fig. 2a**) and skin conductance responses (*F* = 33.436, *p* < 0.001) (**Fig. 2b**), such that participants’ sympathetic drive was significantly greater when anticipating Unpleasant relative to Mild shocks [Contractility: *B* = 0.030 (SE = 0.011), *t* = 2.858 *p* = 0.004; SCRs: *B* = 0.14 (SE = 0.024), *t* = 5.782; *p* < 0.001]. Threat controllability (*p*s > 0.094) did not significantly modulate sympathetic drive, nor did it interact with the modulatory impact of threat unpleasantness (*p*s > 0.234). In summary, anticipating unpleasant shocks—regardless of their controllability—evoked greater cardiac contractility and skin conductance responses than anticipating mild shocks.

**Figure 2.**
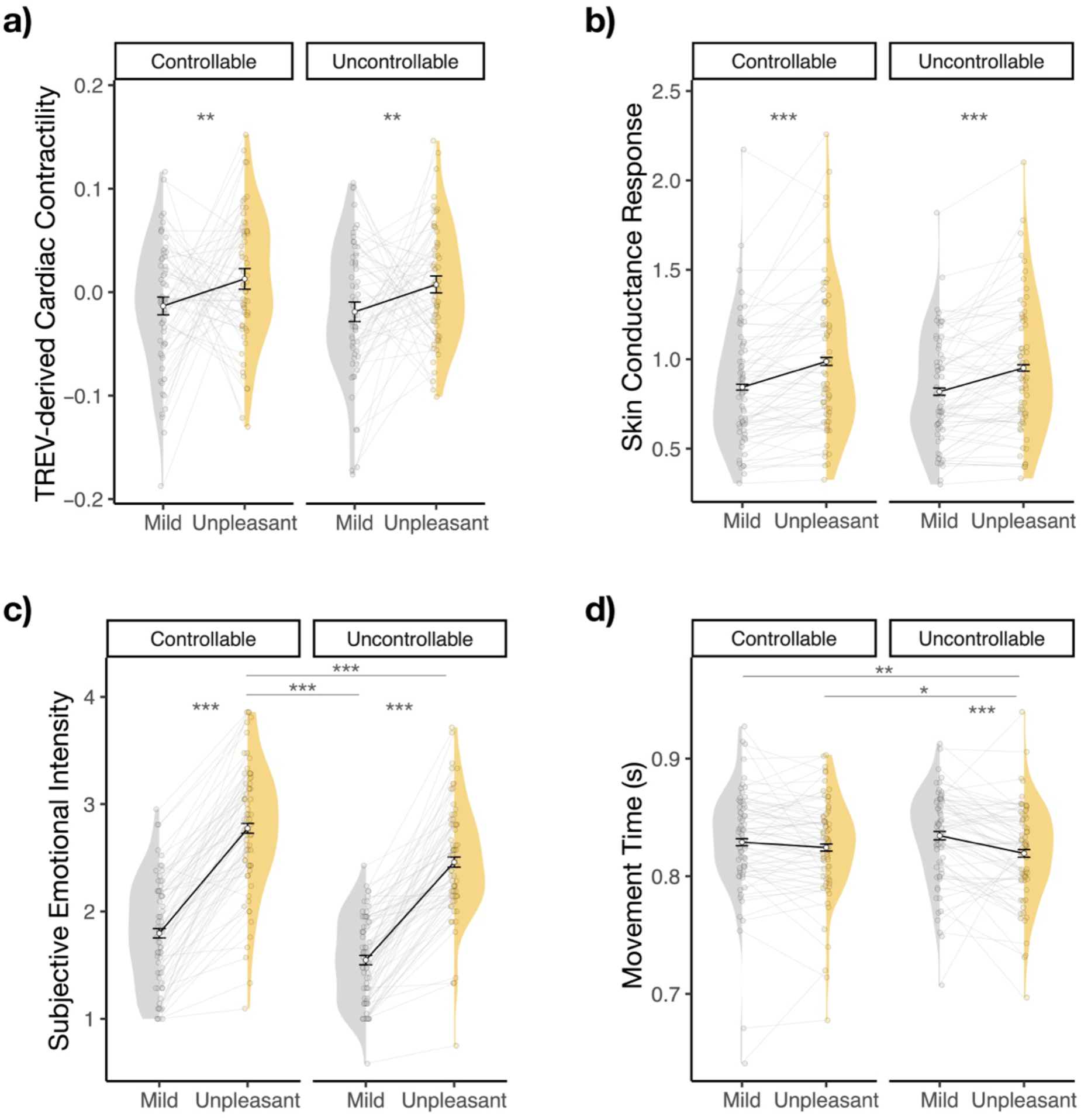
Physiological and Behavioral Modulation During Active Escape. The anticipation of Unpleasant relative to Mild shocks produced larger **a)** cardiac contractility and **b)** skin conductance responses, regardless of threat controllability. **c)** Participants reported experiencing greater emotional intensity during Unpleasant threat anticipation, particularly when anticipating Controllable (*vs*. Uncontrollable) Unpleasant threat. **d)** Participants were quicker to make successful motor responses when anticipating Unpleasant (*vs*. Mild) threat, particularly following Uncontrollable trials. Error bars denote the within-subjects 95% confidence interval for each condition, computed using Morey’s method (Morey, 2008). \**p* < 0.05, \*\**p* < 0.01, \*\*\**p* < 0.001.

When examining participants’ subjective emotional experience during the countdown period, we found that threat unpleasantness robustly increased self-reported emotional intensity (*F* = 283.30, *p* < 0.001), such that participants reported significantly greater emotional intensity when anticipating Unpleasant relative to Mild shocks (*B* = 0.94 (SE = 0.06), *t* = 16.83, *p* < 0.001). This effect was qualified by a significant interaction with controllability (threat unpleasantness * controllability interaction: *F* = 4.457, *p* = 0.035), whereby the anticipation of Controllable Unpleasant shocks elicited higher emotional intensity feelings than all other trial types (*Controllable Unpleasant vs.* Uncontrollable Unpleasant: *B* = 0.318 (SE = 0.050), *t* = 6.318, *p* < 0.001; *vs.* Controllable Mild: *B* = 0.979 (SE = 0.058), *t* = 16.748, *p* < 0.001; *vs.* Uncontrollable Mild: *B* = 1.227 (SE = 0.073), *t* = 16.813, *p* < 0.001) (**Fig. 2c**). Similarly, threat unpleasantness and controllability jointly influenced task performance: Unpleasant (*vs.* Mild) threat anticipation led to faster movement times (*F* = 15.40, *p* < 0.001; *B* = -0.02 (SE = 0.003), *t* = -3.925), particularly during Uncontrollable trials (Threat Unpleasantness * Controllability interaction: *F* = 8.26, *p* = 0.004; Uncontrollable Unpleasant *vs.* Mild: *B* = -0.02 (SE = 0.004), *t* = -4.78, *p* < 0.001; Unpleasant Uncontrollable *vs.* Controllable: *B* = -0.009 (SE = 0.003), *t* = -2.81, *p* = 0.03) (**Fig. 2d**). (See **Figures 2c** and **2d** and Stasiak et al., 2025 for additional details). In summary, anticipating an unpleasant threat powerfully modulated both TREV and skin conductance sympathetic indices, as well as subjective emotional intensity and goal-directed performance, with subjective emotional intensity and task performance additionally modulated by threat controllability.

### TREV-derived contractility is positively associated with electrodermal activity

In light of the robust engagement of both sympathetic channels by threat anticipation, we next examined the relationship between TREV-derived cardiac contractility and SCRs to assess the extent to which these channels reflected overlapping versus unique variance in sympathetic drive. Mixed-effects models revealed a significant, positive association between contractility and SCRs (*F* = 28.581, *p* < 0.001), such that greater cardiac contractility was associated with greater SCR (**Fig. 3**). This positive association was not significantly modulated by task condition (TREV * threat unpleasantness: *p* = 0.316; TREV * controllability: *p =* 0.786; TREV * threat unpleasantness * controllability: *p* = 0.534) (**Supplementary Fig. 1**). Across participants, threat-related changes in TREV and SCR were positively but not significantly associated with one another (Δ Unpleasant − Mild: *r* = 0.224, *p* = 0.089 (**Supplementary Fig. 2**). Taken together, these results provide convergent validity of TREV-derived cardiac contractility as an index of dynamic, within-participant fluctuations in sympathetic drive, while also indicating that TREV and SCRs capture partially distinct components of sympathetic activity.

**Figure 3.**
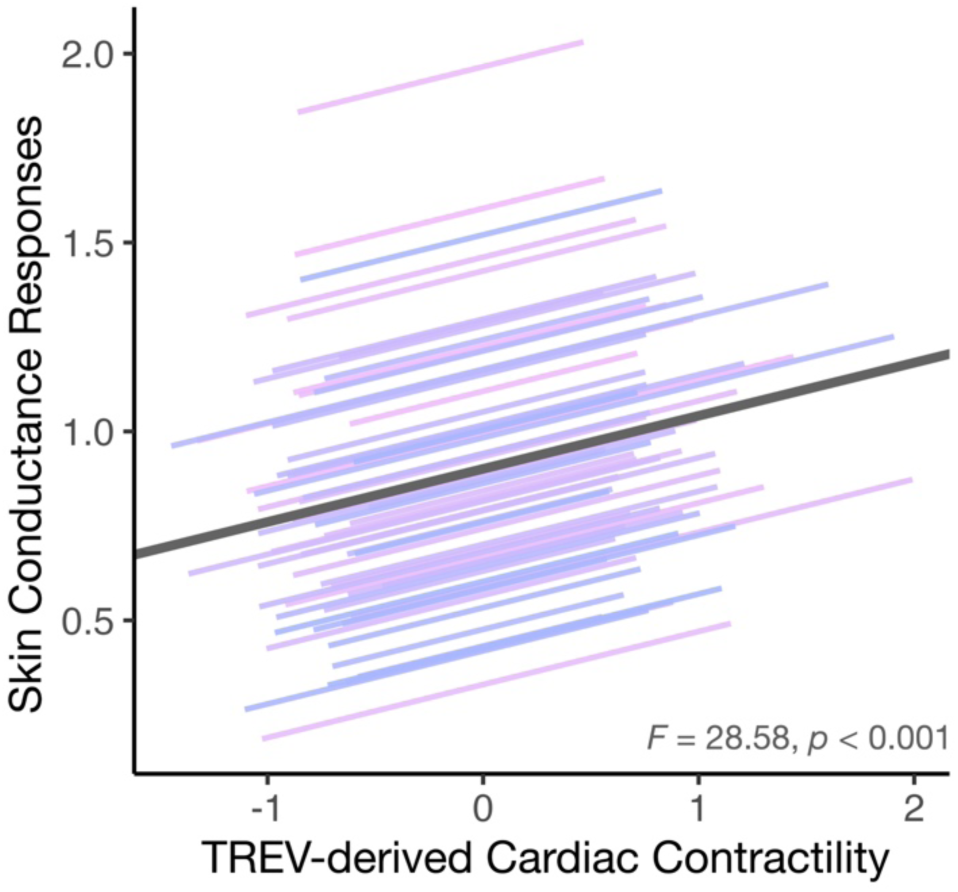
Convergent validity: TREV as a reliable technique for measuring sympathetic drive. TREV-derived cardiac contractility was positively correlated with SCRs, an association that was not modulated by task condition.

### TREV-derived cardiac contractility and skin conductance responses reflect experienced emotional intensity

Prior work suggests weak-to-moderate convergence between sympathetic and experiential response systems (Kreibig, 2010; Mauss et al., 2005). Accordingly, we examined whether participants’ self-reported emotional intensity was associated with TREV-derived cardiac contractility and SCRs, as well as whether these measures tracked overlapping or distinct variance in subjective experience. Mixed-effects models revealed significant positive associations between self-reported emotional intensity and TREV-derived cardiac contractility (*F* = 16.243, *p* < 0.001) (**Fig. 4a**) as well as between emotional intensity and skin conductance responses (*F* = 7.013, *p* = 0.008) (**Fig. 4b**). Critically, post-hoc slope comparisons indicated that the association between sympathetic drive and subjective emotional intensity was significantly stronger for TREV than SCRs (*B* = 0.069 (SE = 0.029), *t* = 2.314, *p* = 0.021). Neither threat unpleasantness nor controllability modulated the association between self-reported emotional intensity and cardiac contractility (*p*s > 0.057) or skin conductance responses (*p*s > 0.19). In summary, the magnitude of sympathetic drive during the anticipation of a potential shock, as indexed by both TREV and SCRs, tracked subsequent ratings of the emotional intensity experienced during the anticipatory period—with TREV showing a significantly stronger association with subjective emotion than SCRs, suggesting partial dissociations between cardiac and electrodermal sympathetic channels.

**Figure 4.**
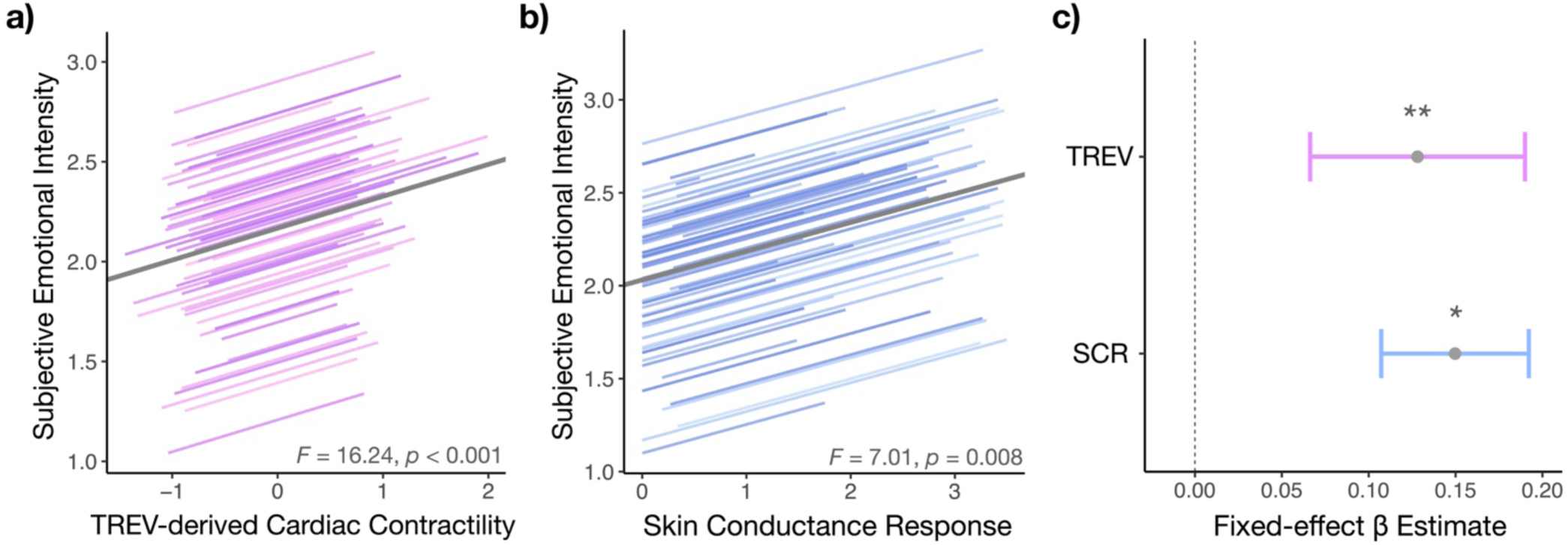
Sympathetic activity tracks self-reported emotional experience. **a)** Participants’ TREV-derived cardiac contractility during the countdown to potential shock administration was positively associated with subjective ratings of emotional intensity provided at the end of each trial. **b)** Participants’ skin conductance responses during the countdown were likewise positively associated with subjective ratings of emotional intensity. **c)** A simultaneous mixed-effects model revealed that TREV and SCRs accounted for unique variance in subjective emotional experience.

To more directly test whether cardiac contractility and skin conductance responses were accounting for shared variance or whether they explained independent variance in self-reported emotional intensity, we ran a simultaneous mixed-effects model entering both cardiac contractility and skin conductance responses as predictors of self-reported emotional intensity ratings. We found that both cardiac contractility and skin conductance responses independently and significantly predicted emotional intensity ratings (TREV: *F* = 10.401, *p* = 0.001; SCRs: *F* = 4.359, *p* = 0.037) (**Fig. 4c**)—further suggesting that TREV and SCRs capture partially separable components of sympathetic arousal that may jointly contribute to subjective emotional experience under threat.

### TREV-derived cardiac contractility uncovers threat-dependent changes in neural engagement

Building on the above-reported threat-related modulation of cardiac contractility and skin conductance responses, and their robust association with subjective emotional experience, we next examined whether anticipatory threat-related increases in sympathetic drive were associated with corresponding changes in neural activation—and whether sympathetic drive and subjective emotional intensity converged on overlapping or separable neural substrates. To this end, we tested whether trial-wise fluctuations in sympathetic drive, indexed by TREV and SCRs, modulated BOLD responses during the threat anticipation countdown period preceding the potential administration of Unpleasant (vs. Mild) shocks using two parametric modulation models (see *Methods* for details). Whole-brain analyses revealed significant clusters of activation correlating with increases in TREV-derived cardiac contractility during Unpleasant (*vs.* Mild) threat anticipation (**Fig. 5**). Specifically, greater trial-wise increases in cardiac contractility during the anticipation of Unpleasant (vs. Mild) shocks were associated with greater activation in the left and medial BA 8 (BA 8B and BA 8m), bilateral posterior intraparietal sulcus (pIPS), as well as in the cerebellum (whole-brain cluster-corrected for multiple comparisons at *Z* > 2.3, *p* < 0.05). Moreover, TREV-modulated activation during threat anticipation was also found in the cerebellar lobule VI, Crus I, and Crus II (**Fig. 5**). Notably, the location of these clusters correspond to regions of the cerebellum previously identified as nodes of the frontoparietal-cerebellar network(Buckner et al., 2011); **Fig. 5** inset).

**Figure 5.**
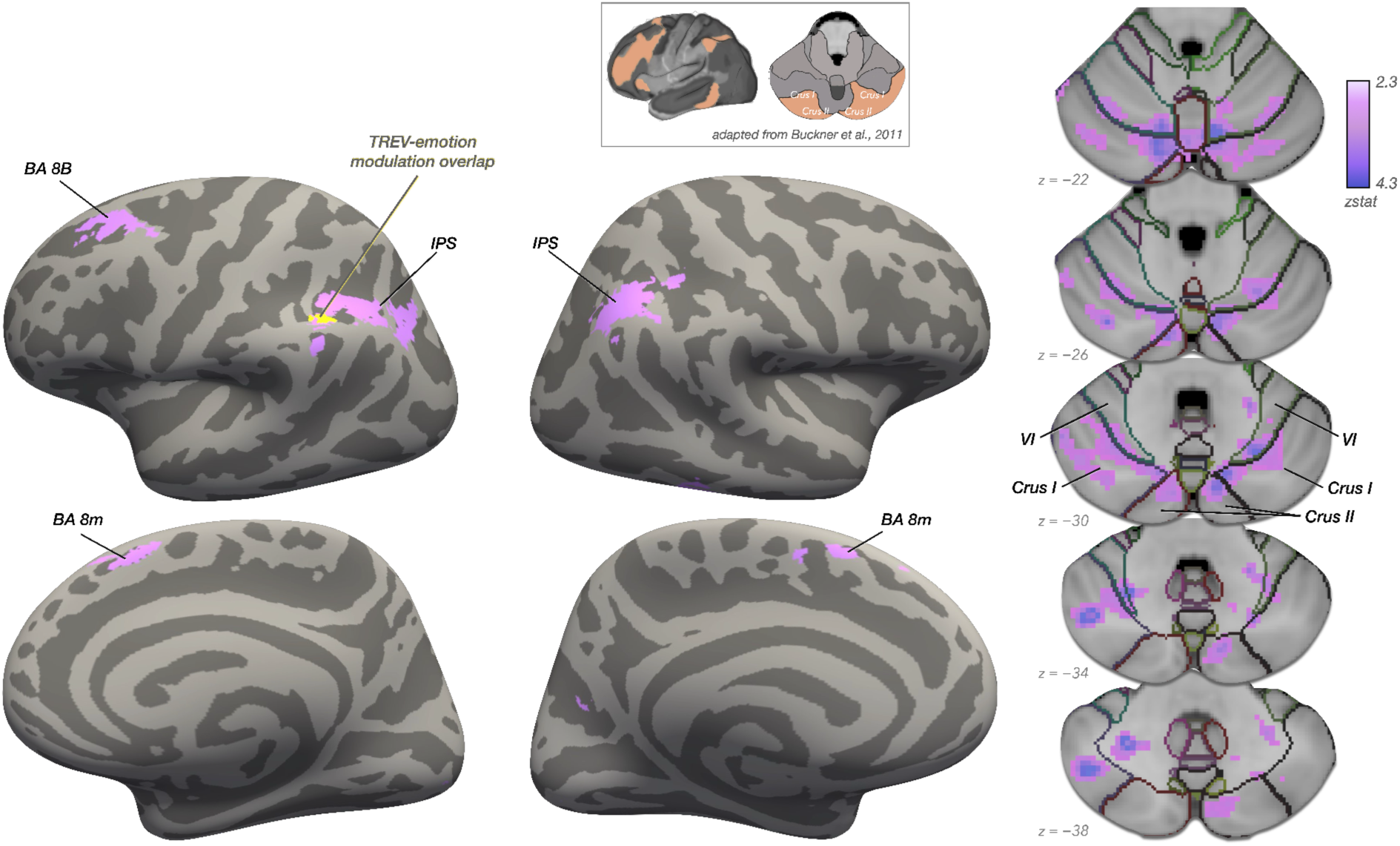
Neural correlates of TREV-derived sympathetic drive during threat anticipation. Greater contractility-related BOLD activation during the anticipation of Unpleasant (vs. Mild) shocks was found in the caudal dorsomedial frontal cortex and posterior parietal cortex, as well as in cerebellar lobule VI, Crus I and Crus II. In the posterior parietal cortex, TREV- and subjective emotional intensity-modulated betas overlapped (shown in yellow). Whole-brain cluster-level corrected for multiple comparisons at *Z* > 2.3, *p* < 0.05.

In contrast, threat-related modulation of BOLD by trial-wise skin conductance responses did not reach significance (Δ Unpleasant − Mild; whole-brain cluster-corrected at *Z* > 2.3, *p* < 0.05; for the unthresholded map, see https://neurovault.org/collections/PEULMFIC/). Moreover, TREV-based BOLD modulation during threat anticipation (Unpleasant vs. Mild) was significantly stronger than that of the SCR-based model (*p* < 0.05, FWE corrected). For additional analyses, including TREV- and SCR-based parametric modulation models collapsed across conditions, see **Supplementary Fig. 3**. Collectively, these findings indicate that changes in TREV-derived cardiac contractility uniquely captures state-dependent changes in neural activation induced by threat anticipation.

Finally, we tested whether changes in TREV-derived sympathetic drive and subjective emotional intensity were associated with overlapping or dissociable neural correlates during Unpleasant (vs. Mild) threat anticipation. To this end, we ran a parametric modulation model of BOLD responses during the anticipation of Unpleasant and Mild shocks as a function of trial-wise variation in subjective ratings of emotional intensity. The resulting map (Δ Unpleasant − Mild) was compared with that produced by the TREV-derived parametric modulation model reported above using a conjunction analysis (see *Methods* for details). Notably, increases in subjective emotional intensity and TREV-derived cardiac contractility during Unpleasant (vs. Mild) threat anticipation were jointly associated with activation in the posterior intraparietal sulcus (whole-brain cluster-corrected for multiple comparisons at *Z* > 2.3, *p* < 0.05) (**Fig. 5**). Together, these findings suggest that TREV-derived cardiac contractility provides a sensitive physiological index of engagement of neural circuitry situated at the intersection of sympathetic modulation and subjective emotional experience.

### TREV-modulated cerebellar activation is associated with motor readiness under threat

To assess the potential functional significance of the neural engagement associated with higher contractility during unpleasant threat anticipation, we next examined whether the magnitude of TREV-modulated neural activation was associated with subsequent motor performance. Given extensive work supporting the cerebellum’s role in motor control (Ebner & Pasalar, 2008; Houk et al., 1997; Mottolese et al., 2013), we correlated the beta estimates (Δ Unpleasant − Mild) extracted from the significant TREV-modulated cerebellar clusters in the parametric model described above (**Fig. 5**) with threat-related changes in movement time (Δ Unpleasant − Mild). We found that across participants, TREV-modulated engagement of cerebellar Crus I and Crus II during unpleasant threat anticipation was negatively associated with movement times (Crus I: r = -0.272, *p* = 0.035; Crus II r = -0.307, *p* = 0.021) (**Fig. 6**). In other words, participants with greater TREV-modulated engagement of cerebellar Crus I and Crus II during unpleasant threat anticipation responded more quickly in these trials—suggesting that neural-cardiac coupling supports adaptive task performance.

**Figure 6.**
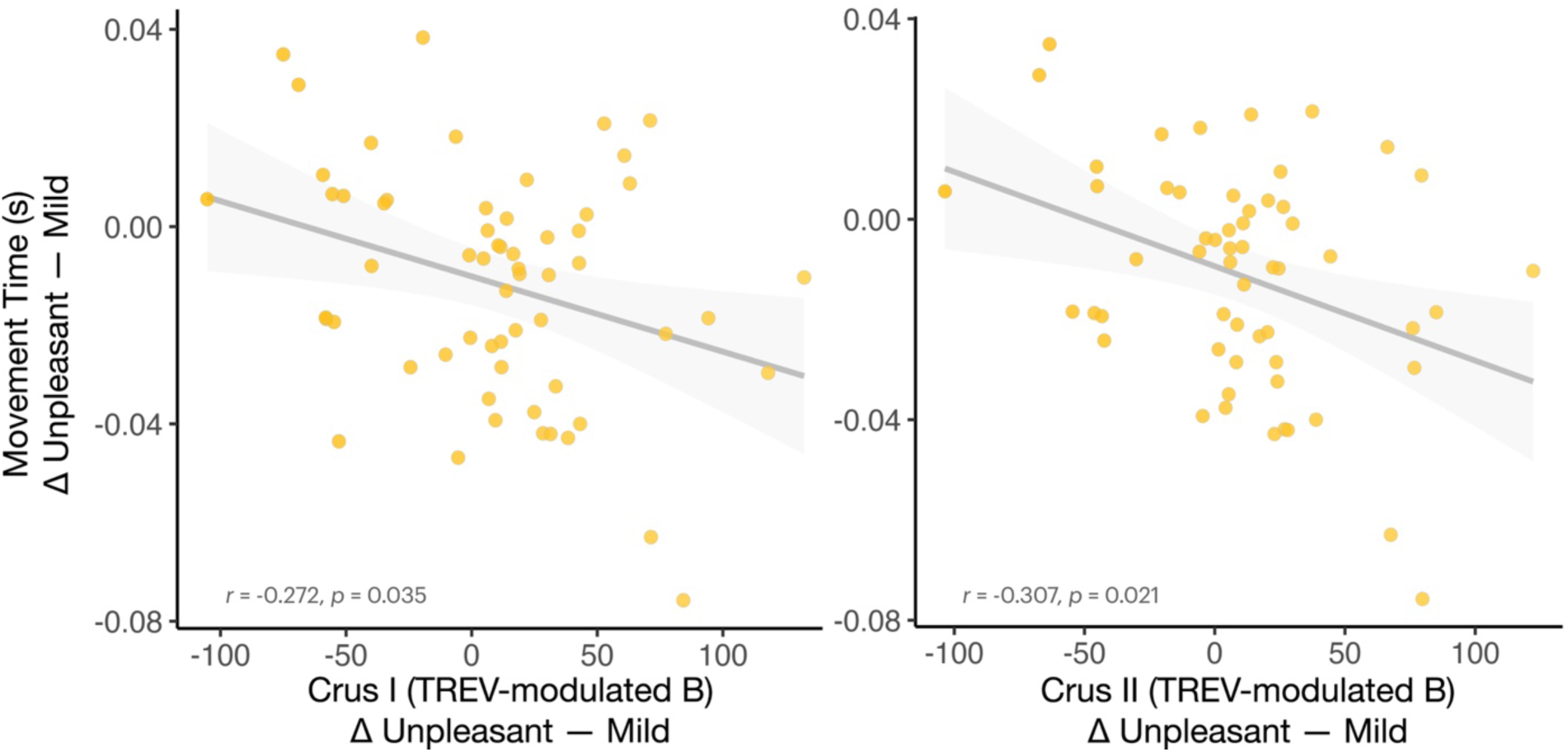
TREV-modulated cerebellar activation facilitates motor responding under threat. Participants with greater contractility-related modulation of activation in cerebellar Crus I (*left*) and Crus II (*right*) during Unpleasant (vs. Mild) threat anticipation showed faster subsequent motor responses following threat.

## Discussion

In the present study, we evaluated TREV-derived cardiac contractility as a physiological index of sympathetic engagement during anticipatory threat-of-shock and tested its relevance for subjective emotional experience, neural activation, and adaptive behavior. Our results demonstrate that TREV provides a sensitive and temporally precise index of sympathetically mediated cardiac contractility that tracks dynamic fluctuations in subjective emotional experience and captures threat-dependent changes in neural engagement. Anticipation of unpleasant threat robustly increased TREV-derived cardiac contractility and SCRs, and these sympathetic indices were positively correlated, providing convergent validity that TREV-derived contractility reflects state-dependent sympathetic drive. Critically, trial-wise fluctuations in TREV-derived cardiac contractility robustly tracked within-subject fluctuations in self-reported emotional intensity and showed stronger associations with emotional experience during threat anticipation than skin conductance responses. Neurally, TREV (but not SCRs) captured threat-related modulation of activation in dorsomedial prefrontal, posterior parietal and cerebellar regions, including cerebellar lobule VI, Crus I, and Crus II, with significantly stronger neural modulation during threat anticipation than that observed for SCRs. Notably, contractility- and subjective emotional-intensity related neural activation converged in posterior intraparietal cortex, identifying a neural substrate linking sympathetic drive to subjective emotional experience. Finally, TREV-modulated cerebellar engagement predicted faster motor responses under threat, suggesting an adaptive cardiac-neural coupling that supports task performance. Collectively, these results position TREV as a novel and sensitive physiological index of sympathetic cardiac drive with high temporal resolution that can be combined with fMRI to advance investigations of autonomic nervous system responding as well as neural-cardiac coupling in response to cognitive, emotional, and social challenges.

First, we provide convergent validity for TREV-derived cardiac contractility as a sympathetic marker by demonstrating reliable coupling with SCRs, consistent with some shared basis of sympathetic output to both skin and muscle. This convergence aligns with empirical accounts of threat anticipation producing behavioral mobilization, modulating sympathetic output to the heart, vasculature, and skeletal musculature (Dampney, 2016; Gianaros & Wager, 2015). Moreover, both TREV-derived cardiac contractility and SCRs robustly tracked within-subject fluctuations in self-reported emotional intensity across trials, in line with prior work linking autonomic responding to subjective affective states (Diemer et al., 2014; Golland et al., 2014; Kreibig, 2010; Stasiak et al., 2023). Importantly, however, each measure explained unique variance in emotional intensity, supporting evidence that physiological and experiential components of emotion show coherent covariance as well as substantial divergence within a shared affective space (Cuve et al., 2023; Eisenbarth et al., 2016; Kragel & Labar, 2013; Kreibig, 2010; Mauss et al., 2005). This divergence aligns with the previously theorized principle of directional fractionation (Berntson et al., 1991; Kreibig, 2010; Lacey, 1967), wherein concurrently activated sympathetic channels express system-specific dynamics depending on behavioral and contextual demands. Together, these findings indicate that TREV and skin conductance responses index partially distinct components of sympathetic influence on affective experience, supporting the coordinated—but non-redundant—contributions of multiple sympathetic effector systems to subjective emotions.

Although both sympathetic channels tracked subjective emotional intensity, TREV and skin conductance responses showed non-overlapping neural correlates of threat-related sympathetic modulation, further supporting a partial dissociation between cardiac and sudomotor efferent responses. Threat-related increases in TREV-derived cardiac contractility, but not SCRs, were associated with increased activation within intraparietal and middle frontal regions, as well as cerebellar lobule VI and Crus I/II—nodes aligning with a frontoparietal-cerebellar network (Buckner et al., 2011; Yeo et al., 2011). The frontoparietal control network is thought to integrate contextual information from the environment with internal representations to guide action (Cole et al., 2014; Vincent et al., 2008) and therefore may closely track threat-related sympathetic changes corresponding to goal-relevant action preparation. Within this network, the posterior intraparietal sulcus was found to be jointly associated with both cardiac contractility and subjective emotional intensity experienced during anticipatory threat. The intraparietal sulcus has reliably been implicated in maintaining body representations (de Jong et al., 2001; Makin et al., 2007; Wolpert et al., 1998), thus potentially supporting a common computational substrate of anticipated threat tied to sensorimotor integration, shared by affective arousal and sympathetic cardiac modulation.

The cerebellum has been increasingly recognized for its role in coordinating predictive motor control with cognitive and affective processes, as well as integrating and modulating somatic and visceral functions (Adamaszek et al., 2017; Sacchetti et al., 2009; Schmahmann & Caplan, 2006; Strick et al., 2009). For instance, emerging evidence has highlighted large scale cortico-cerebellar loops that support cognitive and affective processing (Adamaszek et al., 2017; Buckner et al., 2011; Stoodley & Schmahmann, 2009) in addition to the integration of autonomic signals for adaptive action (Guell et al., 2018; Rudolph et al., 2023). In this context, the contractility-related modulation of cerebellar activation observed here may reflect the integration of sympathetic signals into sensorimotor circuits that coordinate preparatory adaptive behavioral responses. Consistently, contractility-related recruitment of cerebellar Crus I and II during threat anticipation was associated with faster motor responding following unpleasant relative to mild threat anticipation. This finding underscores extant work establishing the cerebellum’s role in fine-tuning adaptive behavior under challenge, consistent with its function in predictive motor control and online sensorimotor optimization (Schmahmann, 1996; Strick et al., 2009). The observed link between contractility-modulated Crus I/II activation and faster motor responding suggests that sympathetic cardiac signals may serve to inform cerebellar computations that support motor readiness—and/or that cerebellar engagement itself modulates sympathetic drive—thereby promoting context-dependent behavioral mobilization (Schmahmann & Caplan, 2006).

Although null results should be interpreted with caution, the partial dissociation observed here between TREV- and skin conductance modulated neural engagement—whereby only TREV, but not SCR, revealed threat-modulated changes in neural responding—suggests that these two indices may capture distinct aspects of sympathetic control. One reason might be methodological: TREV provides finer temporal resolution than skin conductance, which may allow contractility-related fluctuations to align more closely with BOLD hemodynamic changes during threat anticipation. Alternatively, these differences may reflect fundamental distinctions in the neural control architectures of cardiac versus sudomotor sympathetic pathways. Cardiac contractility reflects β-adrenergic sympathetic influences on myocardial vigor via cardiac sympathetic nerves, which directly modulate cardiac output and the capacity to leverage metabolic resources in preparation for action (Dampney, 2016; Levy, 1971; Lymperopoulos et al., 2013). In contrast, skin conductance primarily indexes cholinergic sympathetic innervation of eccrine sweat glands via the sudomotor pathway, a system more closely tied to thermoregulation and generalized arousal than to the dynamic regulation of cardiovascular output required for goal-directed behavior (Boucsein, 2012; Critchley, 2002; Dawson et al., 2007). Such effector-level divergences are paralleled by their partially central neural correlates. For instance, prior neuroimaging studies have linked changes in sympathetic cardiac control to a network encompassing the insula, anterior cingulate, ventromedial prefrontal cortex, and brainstem nuclei (Beissner et al., 2013; Critchley & Harrison, 2013; Dalton et al., 2005; Dundon et al., 2021; Gianaros & Wager, 2015). While skin conductance responses often show modulation by regions that overlap with the network often found in studies of cardiac-related neural correlates—such as the ventromedial prefrontal cortex and anterior cingulate (Boucsein, 2012; Boucsein et al., 2012; Eisenbarth et al., 2016; Nagai et al., 2004; Seifert et al., 2013)—skin conductance responses often also show more diffuse cortical associations and stronger ties to putatively amygdala-originated orienting responses (Phelps et al., 2001; Williams et al., 2001). Consequently, trial-wise variation in cardiac contractility may more closely align with neural systems supporting anticipatory control, action preparation, and effort mobilization under threat, whereas SCRs may primarily reflect more general arousal, and less spatially specific components of sympathetic engagement.

Finally, the present study establishes TREV as a practical and valid tool for simultaneous peripheral-physiological-fMRI acquisition. TREV combines the temporal precision of impedance cardiography with the relative non-obtrusiveness of skin conductance electrode placement, while mitigating respiratory artifacts typical of thoracic measurements that plague ICG metrics (Stump et al., 2024). As mentioned, a critical innovation of TREV is its usage of the derivative of acceleration of ventricular pressure from the obtained impedance wave. This calculation leverages intraventricular pressure as a proxy for contractility, in contrast to PEP calculations which rely on the detection of subtle ICG waveforms (Kosicki et al., 1986; Stump et al., 2024). Collectively, these features make TREV especially suited for experiments requiring continuous, high-resolution measurement of sympathetic drive, such as those examining the neural bases of dynamic shifts in affective states, attention, learning, and cognitive control under stress, and the top-down regulation of arousal.

## Limitations

The findings presented here should be interpreted alongside the following limitations. First, our study focused on negative affect (induced by anticipatory threat), leaving unclear whether TREV-derived cardiac contractility similarly tracks changes in positive affect. Future work would benefit from extending TREV to reward processing paradigms to test whether the observed coupling of contractility with emotional arousal is generalizable across valences. Second, while the present study examined the convergence between TREV and skin conductance, future research should compare the neural and behavioral correlates of TREV with those derived from established measures of cardiac contractility (e.g., ICG/PEP) to evaluate their consistency across methods. Furthermore, it remains unclear whether the observed dissociations between TREV and skin conductance reported here reflect primarily differences in effector systems, their temporal resolution, or both. Finally, while the cerebellar involvement revealed by our analyses suggests functional integration between autonomic control and goal-oriented motor behavior, causal approaches (e.g., perturbation analyses) will be required to confirm mechanistic links between cardiac sympathetic output and adaptive action.

## Conclusion

Together, these findings situate TREV-derived cardiac contractility as a reliable index of sympathetic drive with high temporal resolution that parallels well established autonomic markers while capturing unique aspects of subjective emotional intensity and action mobilization reflected in cardiac-specific neural circuits. By revealing fine-grained links between cardiac dynamics and emotion-relevant neural and behavioral processes, TREV offers a powerful new approach for understanding how the brain and body operate in concert to orchestrate adaptive emotional responding and goal-directed action.

## Data, Materials, and Code Availability

Analyses were run using custom scripts in FSL, Python and R. Python code used for the Active Escape Task is available on GitHub at https://github.com/LEAPNeuroLab/ASAP/tree/main/Active%20Escape. Data and code used for analysis will be made available on Zenodo upon publication of the manuscript.

## Supplementary Information

**Supplementary Figure 1.**
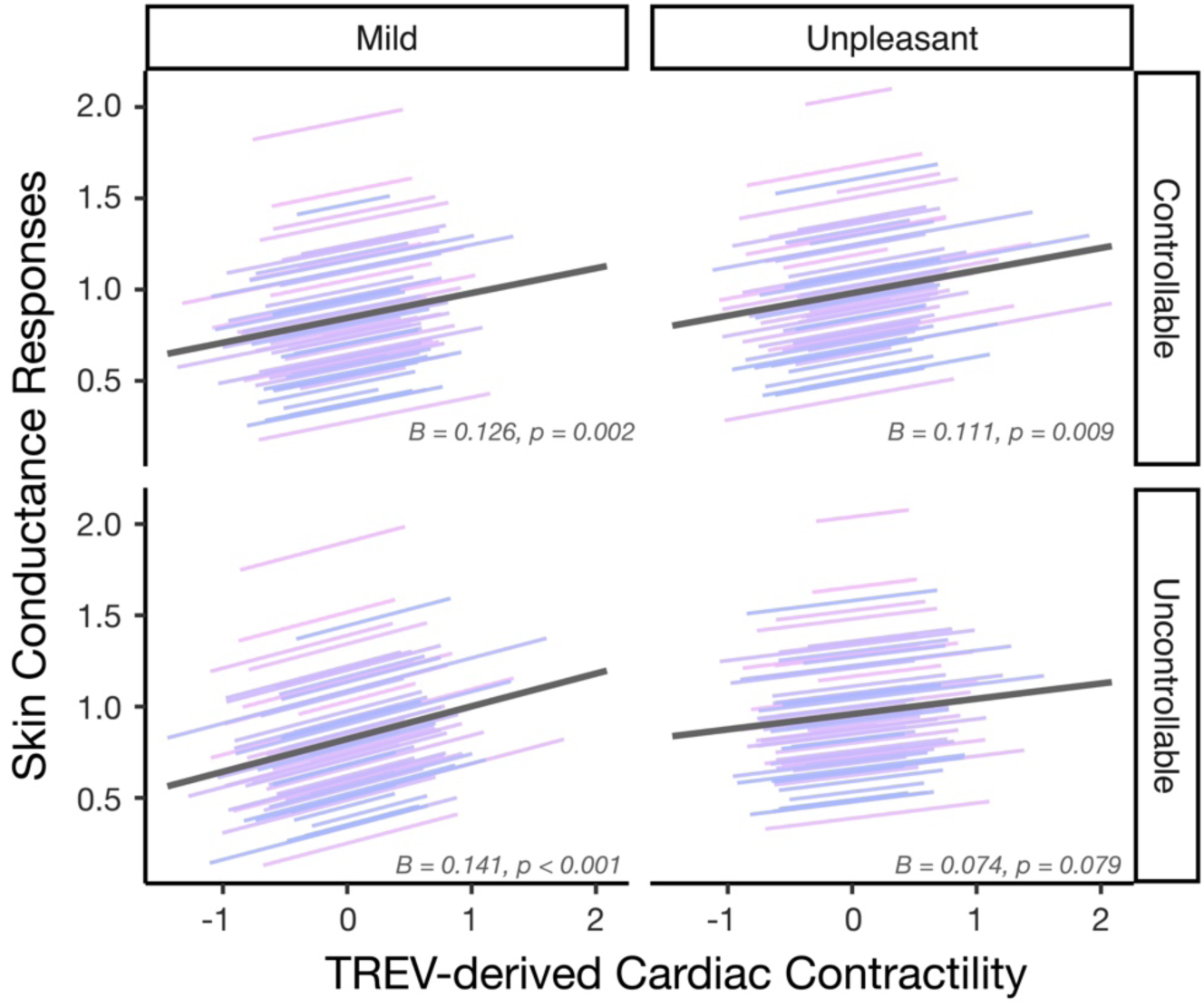
TREV and skin conductance responses plotted by task condition. The observed positive association between cardiac contractility and SCRs (reported in the main text) did not significantly interact with threat unpleasantness nor controllability (TREV * threat unpleasantness: *p* = 0.316; TREV * controllability: *p =* 0.786; TREV * threat unpleasantness * controllability: *p* = 0.534).

**Supplementary Figure 2.**
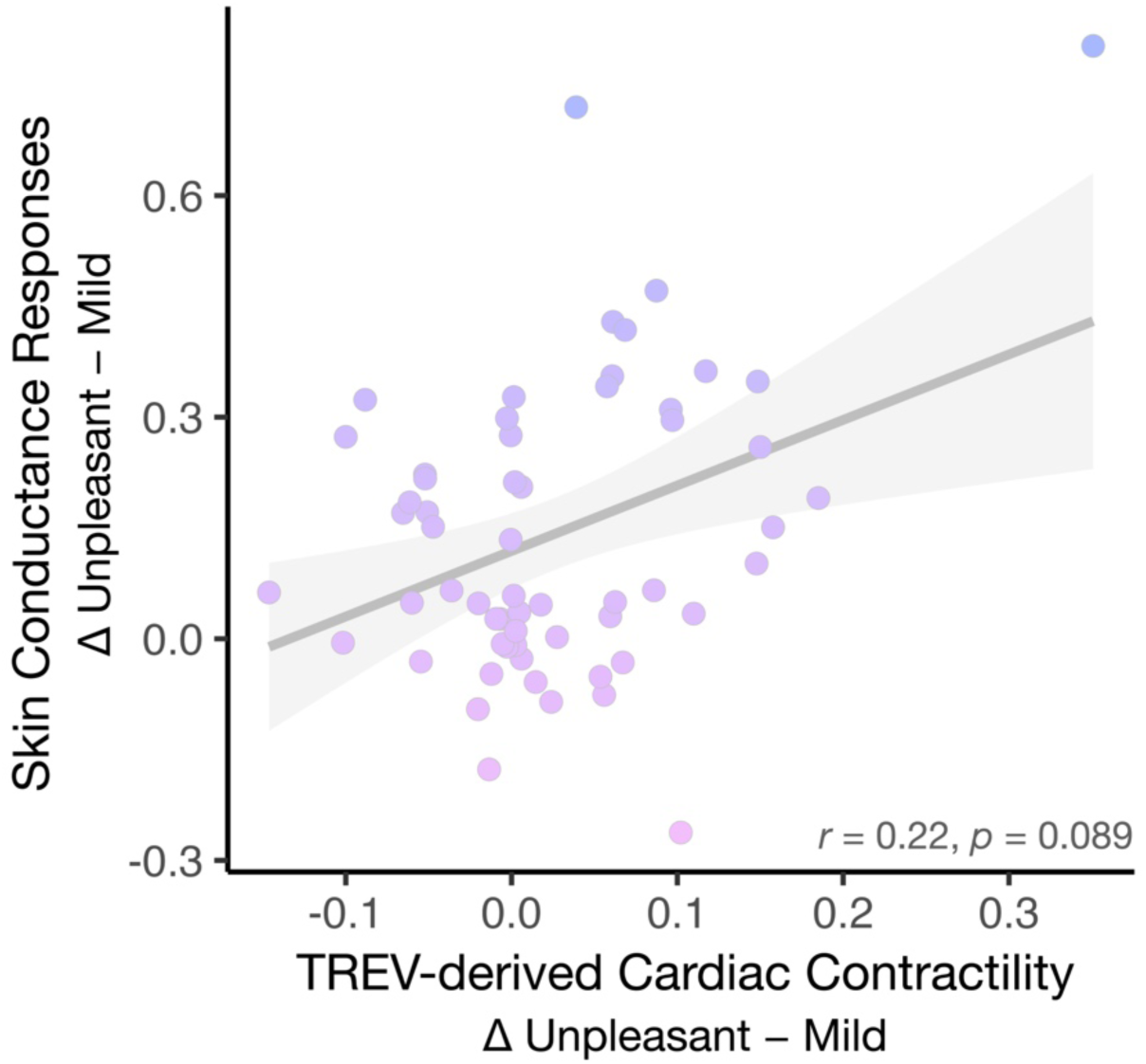
TREV and skin conductance responses across participants. Across participants, threat-related changes in TREV-derived cardiac contractility and SCRs were positively but not significantly correlated (*r* = 0.22, *p* = 0.089).

**Supplementary Figure 3.**
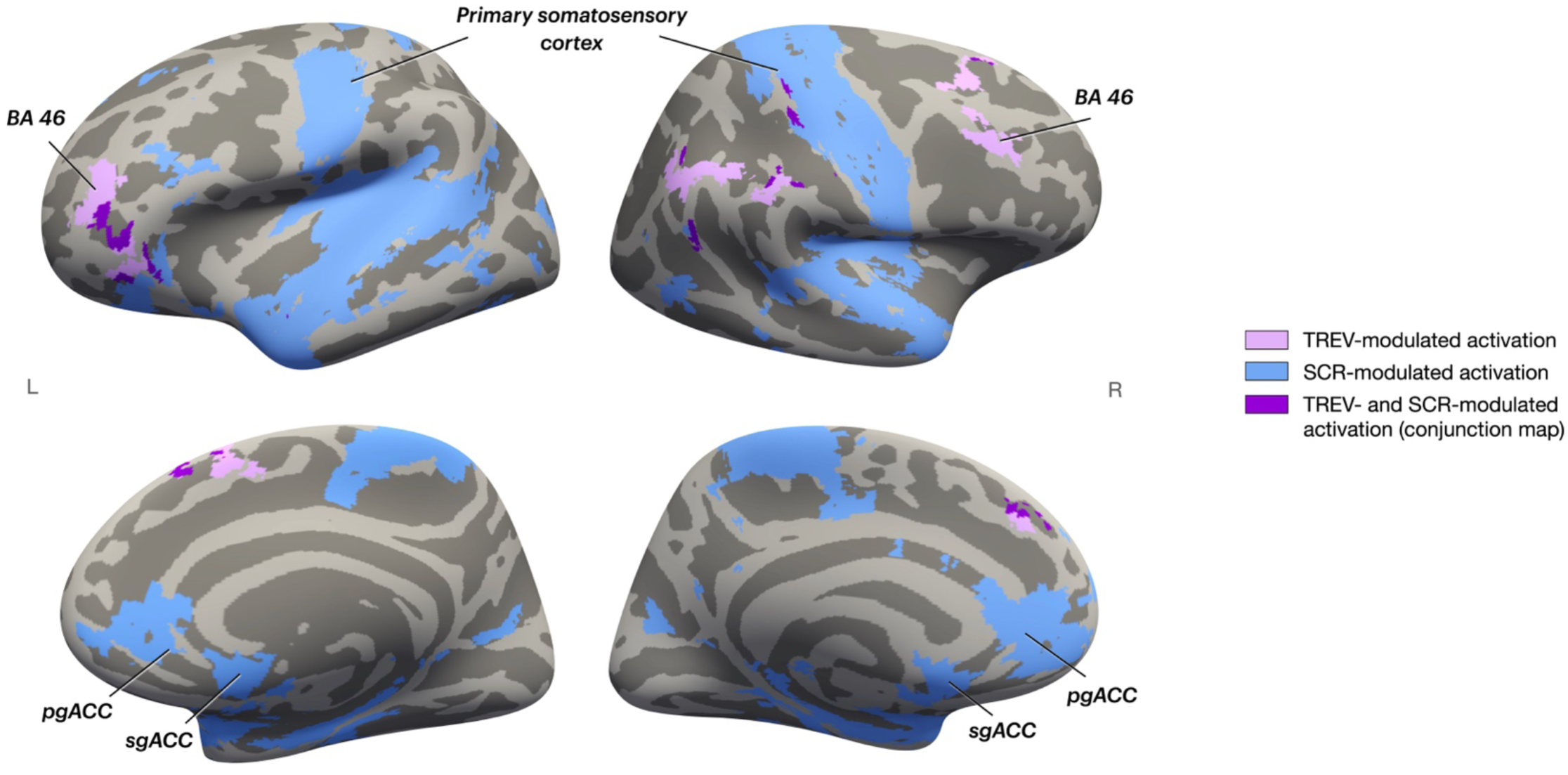
Neural correlates of sympathetic responding. Trial-wise TREV-derived contractility and SCRs were negatively associated with BOLD activation (across task conditions). Significant TREV-modulated betas are shown in lavender, whereas SCR-modulated betas are shown in blue. Clusters of overlapping TREV- and SCR-modulated betas are shown in purple. Whole-brain cluster-level corrected for multiple comparisons at *Z* > 2.3, *p* < 0.05.

## Acknowledgements

This research was funded by NIH grant MH134000 (R.C.L.) and by Aligning Science Across Parkinson’s ASAP-020-519 through the Michael J. Fox Foundation for Parkinson’s Research (S.T.G. and R.C.L.).

## References

Adamaszek, M., D’Agata, F., Ferrucci, R., Habas, C., Keulen, S., Kirkby, K. C., Leggio, M., Mariën, P., Molinari, M., Moulton, E., Orsi, L., Van Overwalle, F., Papadelis, C., Priori, A., Sacchetti, B., Schutter, D. J., Styliadis, C., & Verhoeven, J. (2017). Consensus paper: Cerebellum and emotion. Cerebellum (London, England), 16(2), 552–576.

Adolphs, R., & Andler, D. (2018). Investigating emotions as functional states distinct from feelings. Emotion Review: Journal of the International Society for Research on Emotion, 10(3), 191–201.

Avants, B. B., Tustison, N., & Song, G. (2009). Advanced normalization tools (ANTS). Insight j, 2(365), 1–35.

Bates, D., Mächler, M., Bolker, B., & Walker, S. (2015). Fitting linear mixed-effects models Usinglme4. Journal of Statistical Software, 67(1), 1–48.

Beissner, F., Meissner, K., Bär, K.-J., & Napadow, V. (2013). The autonomic brain: an activation likelihood estimation meta-analysis for central processing of autonomic function. The Journal of Neuroscience: The Official Journal of the Society for Neuroscience, 33(25), 10503–10511.

Benedek, M., & Kaernbach, C. (2010). A continuous measure of phasic electrodermal activity. Journal of Neuroscience Methods, 190(1), 80–91.

Bernstein, D. P. (2009). Impedance cardiography: Pulsatile blood flow and the biophysical and electrodynamic basis for the stroke volume equations. Journal of Electrical Bioimpedance, 1(1), 2–17.

Bernstein, D. P., Henry, I. C., Banet, M. J., & Dittrich, T. (2012). Stroke volume obtained by electrical interrogation of the brachial artery: transbrachial electrical bioimpedance velocimetry. Physiological Measurement, 33(4), 629–649.

Berntson, G. G., Cacioppo, J. T., & Quigley, K. S. (1991). Autonomic determinism: The modes of autonomic control, the doctrine of autonomic space, and the laws of autonomic constraint. Psychological Review, 98(4), 459–487.

Boucsein, W. (2012). Electrodermal activity. https://books.google.com/books?hl=en&lr=&id=6N6rnOEZEEoC&oi=fnd&pg=PR3&dq=boucsein+2012&ots=B4ovsxFJMx&sig=vZSVAb5E750OW6UZtUyNiDah0GU

Boucsein, W., F., Grimnes, S., Ben-Shakhar, G., Roth, W. T., Dawson, M. E., Filion, D. L., & Society for Psychophysiological Research Ad Hoc Committee on Electrodermal Measures. (2012). Publication recommendations for electrodermal measurements: Publication standards for EDA. Psychophysiology, 49(8), 1017–1034.

Bradley, M.M. & Lang, P.J. (2000). Measuring emotion: Behavior, feeling, and physiology. Cognitive Neuroscience of Emotion. https://psycnet.apa.org/record/2000-08961-010

Braithwaite, J. J., Watson, D. G., Jones, R., & Rowe, M. (2015). A guide for analysing electrodermal activity (eda) & skin conductance responses (scrs) for psychological experiments (No. 2). Selective Attention & Awareness Laboratory. https://www.birmingham.ac.uk/documents/college-les/psych/saal/guide-electrodermal-activity.pdf

Büchel, C., Holmes, A., Rees, G., & Friston, K. J. (1998). Characterizing stimulus–response functions using nonlinear regressors in parametric fMRI experiments. NeuroImage, 8, 140–148.

Buckner, R. L., Krienen, F. M., Castellanos, A., Diaz, J. C., & Yeo, B. T. T. (2011). The organization of the human cerebellum estimated by intrinsic functional connectivity. Journal of Neurophysiology, 106(5), 2322–2345.

Buijs, R. M. (2013). The autonomic nervous system: a balancing act. Handbook of Clinical Neurology, 117, 1–11.

Cacioppo, J. T., Tassinary, L. G., & Berntson, G. G. (2000). Psychophysiological science. Handbook of Psychophysiology, 2, 3–23.

Cersosimo, M. G., & Benarroch, E. E. (2013). Central control of autonomic function and involvement in neurodegenerative disorders. Handbook of Clinical Neurology, 117, 45–57.

Chatelain, M., & Gendolla, G. H. E. (2015). Implicit fear and effort-related cardiac response. Biological Psychology, 111, 73–82.

Cieslak, M., Ryan, W. S., Babenko, V., Erro, H., Rathbun, Z. M., Meiring, W., Kelsey, R. M., Blascovich, J., & Grafton, S. T. (2018). Quantifying rapid changes in cardiovascular state with a moving ensemble average. Psychophysiology, 55(4), e13018.

Cieslak, M., Ryan, W. S., Macy, A., Kelsey, R. M., Cornick, J. E., Verket, M., Blascovich, J., & Grafton, S. (2015). Simultaneous acquisition of functional magnetic resonance images and impedance cardiography: ICG during MRI. Psychophysiology, 52(4), 481–488.

Cole, M. W., Repovš, G., & Anticevic, A. (2014). The frontoparietal control system: a central role in mental health: A central role in mental health. The Neuroscientist: A Review Journal Bringing Neurobiology, Neurology and Psychiatry, 20(6), 652–664.

Core Team, R. (2016). R: A language and environment for statistical computing. R Foundation for Statistical Computing, Vienna, Austria. *Http://Www. R-Project. Org/*. https://cir.nii.ac.jp/crid/1574231874043578752

Critchley, H. D. (2002). Electrodermal responses: what happens in the brain. The Neuroscientist: A Review Journal Bringing Neurobiology, Neurology and Psychiatry, 8(2), 132–142.

Critchley, H. D., & Harrison, N. A. (2013). Visceral influences on brain and behavior. Neuron, 77(4), 624–638.

Critchley, H. D., Nagai, Y., Gray, M. A., & Mathias, C. J. (2011). Dissecting axes of autonomic control in humans: Insights from neuroimaging. Autonomic Neuroscience: Basic & Clinical, 161(1–2), 34–42.

Crone, E. A., Somsen, R. J. M., Van Beek, B., & Van Der Molen, M. W. (2004). Heart rate and skin conductance analysis of antecendents and consequences of decision making. Psychophysiology, 41(4), 531–540.

Cuve, H. C. J., Harper, J., Catmur, C., & Bird, G. (2023). Coherence and divergence in autonomic-subjective affective space. Psychophysiology, 60(6), e14262.

Dalton, K. M., Kalin, N. H., Grist, T. M., & Davidson, R. J. (2005). Neural-cardiac coupling in threat-evoked anxiety. Journal of Cognitive Neuroscience, 17(6), 969–980.

Dampney, R. A. (1994). Functional organization of central pathways regulating the cardiovascular system. Physiological Reviews, 74(2), 323–364.

Dampney, R. A. L. (2016). Central neural control of the cardiovascular system: current perspectives. Advances in Physiology Education, 40(3), 283–296.

Dawson, M. E., Schell, A. M., & Filion, D. L. (2007). The electrodermal system. Handbook of Psychophysiology, 2, 200–223.

de Jong, B. M., van der Graaf, F. H., & Paans, A. M. (2001). Brain activation related to the representations of external space and body scheme in visuomotor control. NeuroImage, 14(5), 1128–1135.

Diedrichsen, J., & Zotow, E. (2015). Surface-based display of volume-averaged cerebellar imaging data. PloS One, 10(7), e0133402.

Diemer, J., Mühlberger, A., Pauli, P., & Zwanzger, P. (2014). Virtual reality exposure in anxiety disorders: impact on psychophysiological reactivity. The World Journal of Biological Psychiatry: The Official Journal of the World Federation of Societies of Biological Psychiatry, 15(6), 427–442.

Dum, R. P., Levinthal, D. J., & Strick, P. L. (2016). Motor, cognitive, and affective areas of the cerebral cortex influence the adrenal medulla. Proceedings of the National Academy of Sciences of the United States of America, 113(35), 9922–9927.

Dum, R. P., & Strick, P. L. (2003). An unfolded map of the cerebellar dentate nucleus and its projections to the cerebral cortex. Journal of Neurophysiology, 89(1), 634–639.

Dundon, N. M., Garrett, N., Babenko, V., Cieslak, M., Daw, N. D., & Grafton, S. T. (2020). Sympathetic involvement in time-constrained sequential foraging. Cognitive, Affective & Behavioral Neuroscience, 20(4), 730–745.

Dundon, N. M., Shapiro, A. D., Babenko, V., Okafor, G. N., & Grafton, S. T. (2021). Ventromedial prefrontal cortex activity and sympathetic allostasis during value-based ambivalence. Frontiers in Behavioral Neuroscience, 15, 615796.

Dundon, N. M., Stuber, A., Bullock, T., Garcia, J. O., Babenko, V., Rizor, E., Yang, D., Giesbrecht, B., & Grafton, S. T. (2023). Cardiac-sympathetic contractility and neural alpha-band power: cross-modal collaboration during approach-avoidance conflict. In bioRxiv (Issue 41). bioRxiv. 10.1101/2023.10.10.561785

Ebner, T. J., & Pasalar, S. (2008). Cerebellum predicts the future motor state. Cerebellum (London, England), 7(4), 583–588.

Eisenbarth, H., Chang, L. J., & Wager, T. D. (2016). Multivariate brain prediction of heart rate and skin conductance responses to social threat. The Journal of Neuroscience: The Official Journal of the Society for Neuroscience, 36(47), 11987–11998.

Ganapathy, N., Veeranki, Y. R., & Swaminathan, R. (2020). Convolutional neural network based emotion classification using electrodermal activity signals and time-frequency features. Expert Systems with Applications, 159(113571), 113571.

Gianaros, P. J., & Wager, T. D. (2015). Brain-body pathways linking psychological stress and physical health. Current Directions in Psychological Science, 24(4), 313–321.

Golland, Y., Keissar, K., & Levit-Binnun, N. (2014). Studying the dynamics of autonomic activity during emotional experience: Dynamics of autonomic activity in emotional experience. Psychophysiology, 51(11), 1101–1111.

Guell, X., Schmahmann, J. D., Gabrieli, J. D. E., & Ghosh, S. S. (2018). Functional gradients of the cerebellum. ELife, 7(e36652). 10.7554/eLife.36652

Heinzel, A., Bermpohl, F., Niese, R., Pfennig, A., Pascual-Leone, A., Schlaug, G., & Northoff, G. (2005). How do we modulate our emotions? Parametric fMRI reveals cortical midline structures as regions specifically involved in the processing of emotional valences. Brain Research. Cognitive Brain Research, 25(1), 348–358.

Henderson, L. A., James, C., & Macefield, V. G. (2012). Identification of sites of sympathetic outflow during concurrent recordings of sympathetic nerve activity and fMRI. Anatomical Record (Hoboken, N.J.: 2007), 295(9), 1396–1403.

Henderson, L. A., & Macefield, V. G. (2013). Functional imaging of the human brainstem during somatosensory input and autonomic output. Frontiers in Human Neuroscience, 7, 569.

Henderson, L. A., Stathis, A., James, C., Brown, R., McDonald, S., & Macefield, V. G. (2012). Real-time imaging of cortical areas involved in the generation of increases in skin sympathetic nerve activity when viewing emotionally charged images. NeuroImage, 62(1), 30–40.

Herrald, M. M., & Tomaka, J. (2002). Patterns of emotion-specific appraisal, coping, and cardiovascular reactivity during an ongoing emotional episode. Journal of Personality and Social Psychology, 83(2), 434–450.

Houk, J. C., Buckingham, J. T., & Barto, A. G. (1997). Models of the cerebellum and motor learning. In P. J. Cordo, C. C. Bell, & S. R. Harnad (Eds.), Motor Learning and Synaptic Plasticity in the Cerebellum (pp. 30–45). Cambridge University Press.

Hur, J., Smith, J. F., DeYoung, K. A., Anderson, A. S., Kuang, J., Kim, H. C., Tillman, R. M., Kuhn, M., Fox, A. S., & Shackman, A. J. (2020). Anxiety and the neurobiology of temporally uncertain threat anticipation. The Journal of Neuroscience: The Official Journal of the Society for Neuroscience, 40(41), 7949–7964.

Jansen, A. S., Nguyen, X. V., Karpitskiy, V., Mettenleiter, T. C., & Loewy, A. D. (1995). Central command neurons of the sympathetic nervous system: basis of the fight-or-flight response. Science (New York, N.Y.), 270(5236), 644–646.

Jenkinson, M., Beckmann, C. F., Behrens, T. E. J., Woolrich, M. W., & Smith, S. M. (2012). FSL. NeuroImage, 62(2), 782–790.

Kahle, S., Miller, J. G., Lopez, M., & Hastings, P. D. (2016). Sympathetic recovery from anger is associated with emotion regulation. Journal of Experimental Child Psychology, 142, 359–371.

Karemaker, J. M. (2017). An introduction into autonomic nervous function. Physiological Measurement, 38(5), R89–R118.

Kosicki, J., Chen, L. H., Hobbie, R., Patterson, R., & Ackerman, E. (1986). Contributions to the impedance cardiogram waveform. Annals of Biomedical Engineering, 14(1), 67–80.

Kragel, P. A., & Labar, K. S. (2013). Multivariate pattern classification reveals autonomic and experiential representations of discrete emotions. Emotion (Washington, D.C.), 13(4), 681–690.

Kreibig, S. D. (2010). Autonomic nervous system activity in emotion: a review. Biological Psychology, 84(3), 394–421.

Lacey, J. (1967). Somatic response patterning and stress : some revisions of activation theory. Psychological Stress: Issues in Research. https://cir.nii.ac.jp/crid/1571135650010479232

Lang, P. J. (2014). The motivational organization of emotion: Affect-reflex connections. Emotions. 10.4324/9781315806914-4/motivational-organization-emotion-affect-reflex-connections-peter-lang

Levy, M. N. (1971). Brief reviews: Sympathetic-parasympathetic interactions in the heart. Circulation Research, 29(5), 437–445.

Lymperopoulos, A., Rengo, G., & Koch, W. J. (2013). Adrenergic nervous system in heart failure: pathophysiology and therapy: Pathophysiology and therapy. Circulation Research, 113(6), 739–753.

Makin, T. R., Holmes, N. P., & Zohary, E. (2007). Is that near my hand? Multisensory representation of peripersonal space in human intraparietal sulcus. The Journal of Neuroscience: The Official Journal of the Society for Neuroscience, 27(4), 731–740.

Mauss, I. B., Levenson, R. W., McCarter, L., Wilhelm, F. H., & Gross, J. J. (2005). The tie that binds? Coherence among emotion experience, behavior, and physiology. Emotion (Washington, D.C.), 5(2), 175–190.

Mendes, W. B., Major, B., McCoy, S., & Blascovich, J. (2008). How attributional ambiguity shapes physiological and emotional responses to social rejection and acceptance. Journal of Personality and Social Psychology, 94(2), 278–291.

Morey, R. (2008). Confidence Intervals from Normalized Data: A correction to Cousineau (2005). Tutorials in Quantitative Methods for Psychology, 4, 61–64.

Morrison, S. F. (1999). RVLM and raphe differentially regulate sympathetic outflows to splanchnic and brown adipose tissue. American Journal of Physiology. Regulatory, Integrative and Comparative Physiology, 276(4), R962–R973.

Mottolese, C., Richard, N., Harquel, S., Szathmari, A., Sirigu, A., & Desmurget, M. (2013). Mapping motor representations in the human cerebellum. Brain: A Journal of Neurology, 136(Pt 1), 330–342.

Nagai, Y., Critchley, H. D., Featherstone, E., Trimble, M. R., & Dolan, R. J. (2004). Activity in ventromedial prefrontal cortex covaries with sympathetic skin conductance level: a physiological account of a “default mode” of brain function. NeuroImage, 22(1), 243–251.

Nagel, J. H., Shyu, L. Y., Reddy, S. P., Hurwitz, B. E., McCabe, P. M., & Schneiderman, N. (1989). New signal processing techniques for improved precision of noninvasive impedance cardiography. Annals of Biomedical Engineering, 17(5), 517–534.

Patterson, R. P. (1989). Fundamentals of impedance cardiography. IEEE Engineering in Medicine and Biology Magazine, 8(1), 35–38.

Phelps, E. A., O’Connor, K. J., Gatenby, J. C., Gore, J. C., Grillon, C., & Davis, M. (2001). Activation of the left amygdala to a cognitive representation of fear. Nature Neuroscience, 4(4), 437–441.

Porges, S. W. (1997). Emotion: an evolutionary by-product of the neural regulation of the autonomic nervous system. Annals of the New York Academy of Sciences, 807(1), 62–77.

Rudolph, S., Badura, A., Lutzu, S., Pathak, S. S., Thieme, A., Verpeut, J. L., Wagner, M. J., Yang, Y.-M., & Fioravante, D. (2023). Cognitive-affective functions of the cerebellum. The Journal of Neuroscience: The Official Journal of the Society for Neuroscience, 43(45), 7554–7564.

Sacchetti, B., Scelfo, B., & Strata, P. (2009). Cerebellum and emotional behavior. Neuroscience, 162(3), 756–762.

Scherer, K. R. (2014). On the nature and function of emotion: A component process approach. Approaches to Emotion. 10.4324/9781315798806-18/nature-function-emotion-component-process-approach-klaus-scherer

Schmahmann, J. D. (1996). From movement to thought: anatomic substrates of the cerebellar contribution to cognitive processing. Human Brain Mapping, 4(3), 174–198.

Schmahmann, Jeremy D., & Caplan, D. (2006). Cognition, emotion and the cerebellum [Review of *Cognition, emotion and the cerebellum*]. Brain: A Journal of Neurology, 129(Pt 2), 290–292. Oxford University Press (OUP).

Seifert, F., Schuberth, N., De Col, R., Peltz, E., Nickel, F. T., & Maihöfner, C. (2013). Brain activity during sympathetic response in anticipation and experience of pain. Human Brain Mapping, 34(8), 1768–1782.

Sherwood, A., Allen, M. T., Fahrenberg, J., Kelsey, R. M., Lovallo, W. R., & van Doornen, L. J. (1990). Methodological guidelines for impedance cardiography. Psychophysiology, 27(1), 1–23.

Sinha, R., Lovallo, W., & Parsons, O. (1992). Cardiovascular differentiation of emotions. Psychosomatic Medicine, 54, 422–435.

Smith, S. M., Jenkinson, M., Woolrich, M. W., Beckmann, C. F., Behrens, T. E. J., Johansen-Berg, H., Bannister, P. R., De Luca, M., Drobnjak, I., Flitney, D. E., Niazy, R. K., Saunders, J., Vickers, J., Zhang, Y., De Stefano, N., Brady, J. M., & Matthews, P. M. (2004). Advances in functional and structural MR image analysis and implementation as FSL. NeuroImage, 23 *Suppl 1*, S208–19.

Stasiak, J. E., Mitchell, W. J., Reisman, S. S., Gregory, D. F., Murty, V. P., & Helion, C. (2023). Physiological arousal guides situational appraisals and metacognitive recall for naturalistic experiences. Neuropsychologia, 180(108467), 108467.

Stasiak, J. E., Wang, J., Dundon, N. M., Rizor, E. J., Villanueva, C. M., Barandon, P. L., Grafton, S. T., & Lapate, R. C. (2025). Integrated representations of threat and controllability in the lateral frontal pole. Journal of Cognitive Neuroscience, 1–16.

Stoodley, C. J., & Schmahmann, J. D. (2009). Functional topography in the human cerebellum: a meta-analysis of neuroimaging studies. NeuroImage, 44(2), 489–501.

Strick, P. L., Dum, R. P., & Fiez, J. A. (2009). Cerebellum and nonmotor function. Annual Review of Neuroscience, 32(1), 413–434.

Stump, A., Gregory, C., Babenko, V., Rizor, E., Bullock, T., Macy, A., Giesbrecht, B., Grafton, S. T., & Dundon, N. M. (2024). Non-invasive monitoring of cardiac contractility: Trans-radial electrical bioimpedance velocimetry (TREV). Psychophysiology, 61(1), e14411.

Venables, P. H., & Mitchell, D. A. (1996). The effects of age, sex and time of testing on skin conductance activity. Biological Psychology, 43(2), 87–101.

Vincent, J. L., Kahn, I., Snyder, A. Z., Raichle, M. E., & Buckner, R. L. (2008). Evidence for a frontoparietal control system revealed by intrinsic functional connectivity. Journal of Neurophysiology, 100(6), 3328–3342.

Wehrwein, E. A., Orer, H. S., & Barman, S. M. (2016). Overview of the anatomy, physiology, and pharmacology of the autonomic nervous system. Comprehensive Physiology, 6(3), 1239–1278.

Williams, L. M., Phillips, M. L., Brammer, M. J., Skerrett, D., Lagopoulos, J., Rennie, C., Bahramali, H., Olivieri, G., David, A. S., Peduto, A., & Gordon, E. (2001). Arousal dissociates amygdala and hippocampal fear responses: evidence from simultaneous fMRI and skin conductance recording. NeuroImage, 14(5), 1070–1079.

Winkler, A. M., Ridgway, G. R., Webster, M. A., Smith, S. M., & Nichols, T. E. (2014). Permutation inference for the general linear model. Neuroimage, 92, 381–397.

Wolpert, D. M., Goodbody, S. J., & Husain, M. (1998). Maintaining internal representations: the role of the human superior parietal lobe. Nature Neuroscience, 1(6), 529–533.

Woltjer, H. H., Bogaard, H. J., & de Vries, P. M. (1997). The technique of impedance cardiography. European Heart Journal, 18(9), 1396–1403.

Yeo, B. T. T., Krienen, F. M., Sepulcre, J., Sabuncu, M. R., Lashkari, D., Hollinshead, M., Roffman, J. L., Smoller, J. W., Zöllei, L., Polimeni, J. R., Fischl, B., Liu, H., & Buckner, R. L. (2011). The organization of the human cerebral cortex estimated by intrinsic functional connectivity. Journal of Neurophysiology, 106(3), 1125–1165.

Ziemssen, T., & Siepmann, T. (2019). The investigation of the cardiovascular and sudomotor autonomic nervous system-A review. Frontiers in Neurology, 10, 53.

